# *Phytophthora sojae* effector Avr1d functions as E2 competitor and inhibits ubiquitination activity of GmPUB13 to facilitate infection

**DOI:** 10.1101/2020.09.19.304535

**Authors:** Yachun Lin, Qinli Hu, Jia Zhou, Weixiao Yin, Deqiang Yao, Yuanyuan Shao, Yao Zhao, Baodian Guo, Yeqiang Xia, Qian Chen, Yan Wang, Wenwu Ye, Qi Xie, Brett M. Tyler, Weiman Xing, Yuanchao Wang

## Abstract

Oomycete pathogens such as *Phytophthora* secrete a repertoire of effectors to host cells to manipulate host immunity and benefit infection. In this study, we found that an RxLR effector, Avr1d, promoted *Phytophthora sojae* infection in soybean hairy-roots. Using a yeast two-hybrid screen, we identified the soybean E3 ubiquitin ligase GmPUB13 as a host target for Avr1d. By co-immunoprecipitation, gel infiltration and ITC assays, we confirmed that Avr1d interacts with GmPUB13 both *in vivo* and *in vitro*. Furthermore, we found that Avr1d inhibits the E3 ligase activity of GmPUB13. The crystal structure of Avr1d in complex with GmPUB13 was solved and revealed that Avr1d occupies the binding site for E2 ubiquitin conjugating enzyme on GmPUB13. In line with this, Avr1d competed with E2 ubiquitin conjugating enzymes for GmPUB13 binding *in vitro,* thereby decreasing the E3 ligase activity of GmPUB13. Meanwhile, we found that inactivation of the ubiquitin ligase activity of GmPUB13 stabilized GmPUB13 by blocking GmPUB13 degradation. Silencing of GmPUB13 in soybean hairy-roots decreased *P. sojae* infection, suggesting that GmPUB13 acts as a susceptibility factor, negatively regulating soybean resistance against *P. sojae*. Altogether, this study highlights a novel virulence mechanism of *Phytophthora* effectors, by which Avr1d competes with E2 for GmPUB13 binding to repress the GmPUB13 E3 ligase activity and thereby stabilizing the susceptibility factor GmPUB13 to facilitate *Phytophthora* infection. This is the first study to unravel the structural basis for modulation of host targets by *Phytophthora* effectors and will be instrumental for boosting plant resistance breeding.

**Significance Statement:** Ubiquitination acts as a crucial regulator in plant immunity. Accordingly, microbial pathogens secrete effectors to hijak host ubiquitination system. However, the molecular mechanisms by which microbial effectors modulate host ubiquitination system are not yet clear. Here, we found that the *Phytophthora sojae* effector Avr1d physically binds to the U-box type E3 ligase GmPUB13, a susceptibility factor in soybean. The crystal structure of Avr1d in complex with GmPUB13 revealed that Avr1d occupies the binding site in GmPUB13 for the E2 ubiquitin conjugating enzyme and competes with E2 for physical binding to GmPUB13. Avr1d stabilized GmPUB13 by suppressing the self-ubiquitination activity of GmPUB13 and thereby promoting *Phytophthora* infection. This study provides structural basis for modulation of host targets by *Phytophthora* effectors.

## Introduction

In nature, plants are continuously challenged by various microbes. During long-term co-evolution, plants have developed innate immune systems to discriminate among microbes by recognizing Microbe-Associated Molecular Patterns (MAMP) and microbial effectors, and thereby mounting MAMP/PAMP-Triggered Immunity (MTI/PTI) and Effector-Triggered Immunity (ETI) responses, respectively. For productive infection, plant pathogens secrete a complex repertoire of effectors to modulate host immunity to favor infection (1).

Ubiquitination has emerged as a crucial posttranslational modification of proteins, as it plays an important role in regulating various cellular processes, including plant immune responses (2, 3). In general, ubiquitination involves addition of ubiquitin to target proteins by sequential actions of an E1 ubiquitin activating enzyme, an E2 ubiquitin conjugating enzyme and an E3 ubiquitin ligase. Ubiquitination can trigger degradation or may modify the function of the target protein. E3 ubiquitin ligases AtPUB13 and AtPUB12 are well-studied negative regulators of *A. thaliana* immunity, which attenuate the immune responses triggered by the bacteria PAMP flg22 (4, 5). AtPUB13 also ubiquitinates the chitin receptor AtLYK5, which is required for AtLYK5 turnover; the mutant *pub13* is more sensitive to chitin treatment (6).

The oomycete pathogen *P. sojae* is the causal agent of root and stem rot of soybean, which accounts for millions of dollars of annual loss worldwide (7). During infection, *P. sojae* secretes a repertoire of effectors with a conserved N-terminal ‘RxLR’ motif into host cells (8). Avr1d (Avh6) is one such effector and previously was identified as an avirulence effector which can be recognized by soybean plants carrying the resistance gene *Rps1d,* triggering ETI (9). In this study, we found that Avr1d was a virulence factor that promotes *P. sojae* infection when overexpressed in soybean hairy-roots. During *P. sojae* infection, Avr1d targets two highly homologous U-box ARM-repeats type E3 ligases in soybean, Glyma12g06860 and Glyma11g14910, which are phylogenetically close to AtPUB13 and were named as GmPUB13 and GmPUB13-like (GmPUB13L), respectively. Silencing GmPUB13 and GmPUB13L in soybean hairy-roots increased the resistance to *P. sojae*. Avr1d physically binds to GmPUB13 and blocks the ubiquitin ligase activity of GmPUB13 by competing with E2 ubiquitin conjugating enzyme for the U-box domain. This study reveals the structural basis for modulation of host targets by *Phytophthora* effectors and provides mechanistic insight into the suppression of host immunity by *Phytophthora* effectors.

## Results

### Avr1d promotes *P. sojae* infection

In our previous study, we found Avr1d could be recognized by soybean isogenic lines carrying *Rps1d* (9). To figure out whether Avr1d acts as a virulence effector, we overexpressed Avr1d (without signal peptide) fused with an N-terminal enhanced green fluorescence protein (eGFP) in soybean hairy-roots and performed infection assays using mRFP-labeled *P. sojae*. *P. sojae* infection was evaluated by quantifying the number of produced oospores and *P. sojae* biomass at 2 days post inoculation. Both the produced oospores and biomass of *P. sojae* were much greater in the hairy-roots overexpressing eGFP-Avr1d than in the eGFP control (Fig. 1*A*). These results suggested that Avr1d could promote *P. sojae* infection

**Fig. 1.**
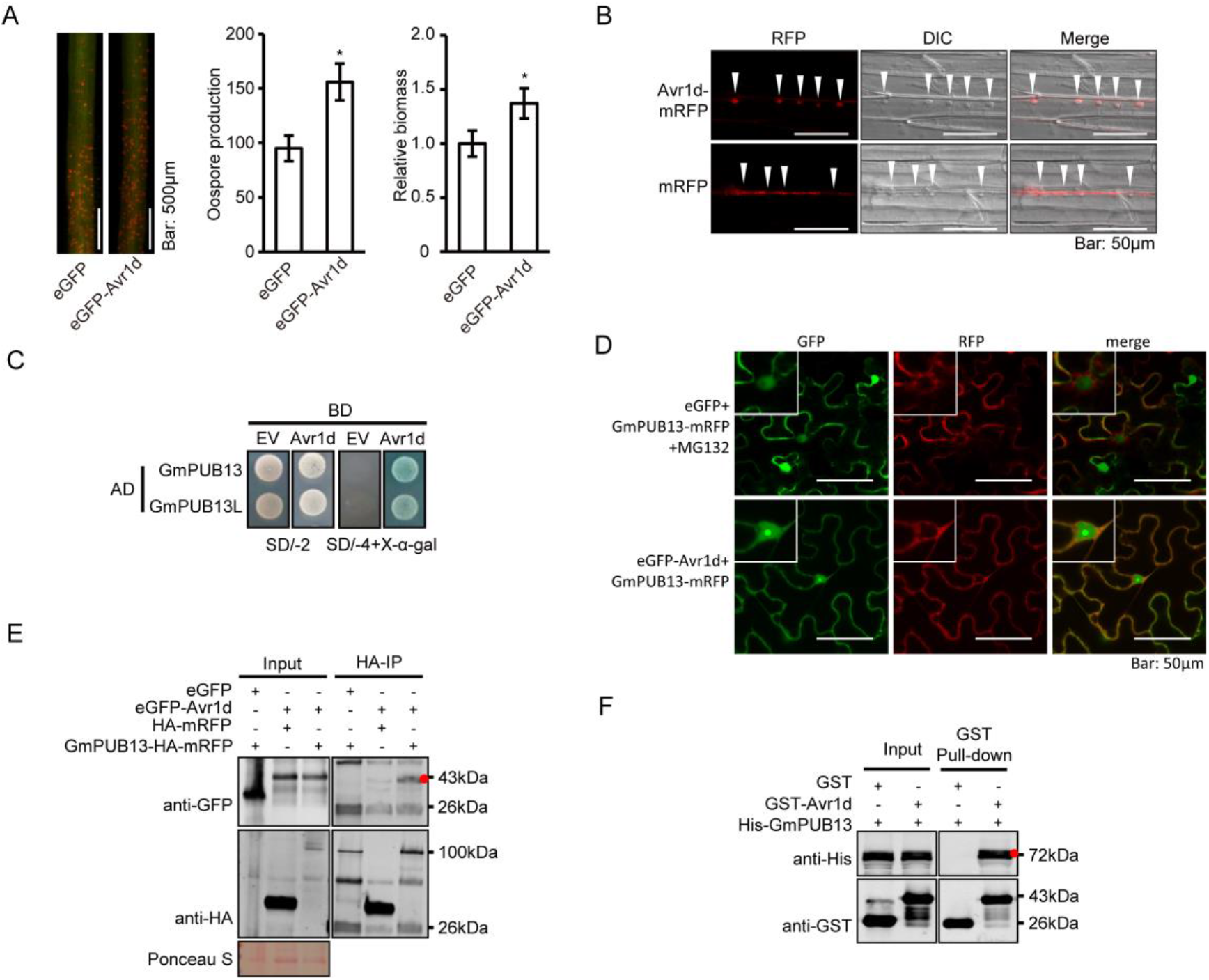
Avr1d promotes *P. sojae* infection and targets to GmPUB13. **(*A*)** Overexpression of Avr1d in soybean hairy-roots promotes *P. sojae* infection. The fluorescent hairy-roots expressing eGFP or eGFP-Avr1d were inoculated with mRFP-labeled *P. sojae* hyphae plugs on the tips (left). Scale bar = 500 μm. The oospore production was observed under the fluorescence microscopy (middle). Relative biomass of *P. sojae* was determined by qPCR at 48 hours post inoculation (hpi) (right). Asterisks indicated t-test p < 0.05. These experiments were repeated 3 times with similar results. **(*B*)** Avr1d is secreted through haustoria. The etiolated soybean seedlings were inoculated with *P. sojae* zoospore expressing Avr1d-mRFP or mRFP. Triangles indicated *P. sojae* haustoria. Scale bar = 50 μm. **(*C*)** The interaction of Avr1d with GmPUB13 and GmPUB13L in yeast. Avr1d interacted with GmPUB13 and GmPUB13L, as indicated by yeast two-hybrid assays. EV, empty vector; SD/-2 medium: SD/-Leu/-Trp medium; SD/-4 + X-α-gal: SD/-Ade/-His/-Leu/-Trp + X-α-gal medium. **(*D*)** eGFP-Avr1d and GmPUB13-HA-mRFP co-localized at cytoplasm in *N. benthamiana* cells. eGFP and GmPUB13-HA-mRFP were co-expressed in *N. benthamiana* leaves (infiltrated with 50 μM MG132 12 hours before observation) by agroinfiltration. eGFP-Avr1d and GmPUB13-HA-mRFP were co-expressed in *N. benthamiana* leaves by agroinfiltration. The leaves were observed under confocal microscopy 2 days post infiltration. The smaller pictures on the upper left were a bigger view of nucleus region in the main picture. Scale bar: 50 μm. **(*E*)** Confirmation of the interaction between Avr1d and GmPUB13 by Co-IP. Pairs of eGFP and GmPUB13-HA-mRFP, eGFP-Avr1d and HA-mRFP, eGFP-Avr1d and GmPUB13-HA-mRFP were co-expressed in *N. benthamiana* leaves, respectively. Total protein extractions of the leaves were incubated with agarose beads conjugated anti-HA mouse monoclonal antibody followed by washing with PBS buffer for 5 times. Proteins in each sample were detected by western blot with anti-GFP and anti-HA antibodies. Red dot indicated the protein size of eGFP-Avr1d. Ponceau S: ponceau staining indicates the rubisco protein. Molecular mass markers are shown (kDa). **(*F*)** Confirmation of the interaction between Avr1d and GmPUB13 by GST pull down. Protein extracts of GST-Avr1d and His-GmPUB13 or GST-tag and His-GmPUB13 were mixed and incubated with glutathione sepharose 4B beads respectively, and followed by washing with PBS buffer for 5 times. Proteins in the input samples and pull down samples were detected by western blot with anti-His and anti-GST antibodies. Red dot indicated the protein size of His-GmPUB13. Molecular mass markers are shown (kDa).

To confirm secretion of Avr1d by *P. sojae*, Avr1d with native signal peptide fused with monomeric red fluorescence protein (mRFP) at the C-terminus was overexpressed in the wild type *P. sojae* strain P6497. The mRFP-labeled *P. sojae* was used as a control (10). Protein expression was confirmed by western blot with anti-mRFP (*SI Appendix,* Fig. S1). Confocal microscopy visualization of infected hypocotyl epidermal cells showed that the red fluorescence of Avr1d-mRFP concentrated specifically at the haustoria, while the control mRFP localized in the cytosol, without significant concentration in the haustoria (Fig. 1*B*). This result suggested that the haustoria were a primary site of Avr1d secretion during infection.

### Avr1d physically interacts with GmPUB13 *in vitro* and *in vivo*

To uncover the molecular mechanism underlying the function of Avr1d in promoting *P. sojae* infection, we performed yeast two-hybrid screening to identify potential soybean targets of Avr1d. Through screening, we identified two highly homologous genes, Glyma12g06860 and Glyma11g14910 that encode U-box ARM-repeats-type E3 ligases. Phylogenetic analysis indicated that these two genes were homologous to AtPUB13 of *Arabidopsis thaliana* (*SI Appendix,* Fig. S2). Glyma12g06860 and Glyma11g14910 shared an amino acid identity of 97% and we named them GmPUB13 and GmPUB13L, respectively. Further, yeast two-hybrid assays supported that Avr1d interacted with both GmPUB13 and GmPUB13L (Fig. 1*C*). In support of the interaction between Avr1d and GmPUB13, we co-expressed Avr1d and GmPUB13 in *Nicotiana benthamiana* and found eGFP-Avr1d and GmPUB13-HA-mRFP co-localized to the cytoplasm under the confocal fluorescence microscopy (Fig. 1*D*). Furthermore, GmPUB13-HA-mRFP co-immunoprecipitated from *N. benthamiana* cell extracts with eGFP-Avr1d, but not with eGFP (Fig. 1*E*). To determine whether Avr1d physically interacted with GmPUB13 *in vitro*, we performed pull-down assays and found His-GmPUB13 could be pulled down by GST-Avr1d, but not by GST (Fig. 1*F*). The physical interaction between Avr1d and GmPUB13 was further confirmed by gel filtration chromatography (*SI Appendix,* Fig. S3). Together, these data demonstrated that Avr1d physically interacted with GmPUB13 both *in vivo* and *in vitro*.

### Crystal structure of the complex of Avr1d with the GmPUB13 U-box domain

Avr1d belongs to the class of RxLR effectors that carry a signal peptide followed by the RxLR motif and an effector domain (*SI Appendix,* Fig. S4*A*) (9). To characterize the molecular mechanism by which Avr1d contributes to the virulence of *P. sojae*, we ascertained the structure of the effector domain of Avr1d complexed with the U-box domain of GmPUB13 (Fig. 2*A*, *SI Appendix,* Fig. S4*A*). The structure of the complex was determined at 2.7 Å resolution using single wavelength anomalous diffraction. The model was refined to final R_work_ and R_free_ values of 23.5% and 24.9%, respectively (*SI Appendix,* Table S1). The asymmetrical unit of the crystals contained one Avr1d-GmPUB13 U-box complex (PDB: 7C96).

**Fig. 2.**
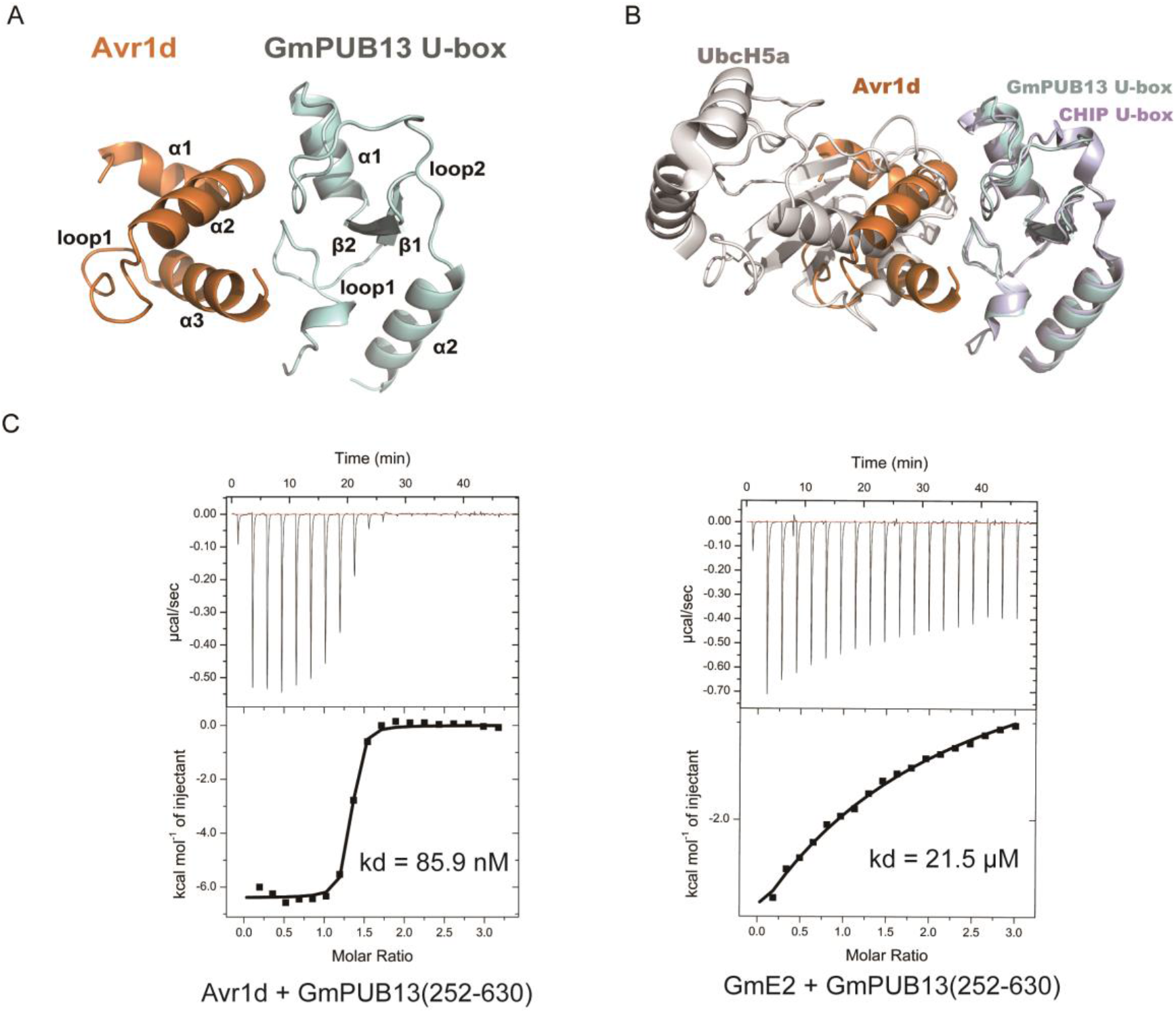
Avr1d interacts with GmPUB13 by occupying E2 binding site of GmPUB13. **(*A*)** Overall structure of Avr1d-GmPUB13 U-box. Avr1d (orange) and GmPUB13 U-box (cyan) are shown in cartoon form. **(*B*)** Superposition of crystal structures of Avr1d-GmPUB13 U-box and UbcH5a-CHIP U-box (PDB: 2OXQ). Avr1d (orange), GmPUB13 U-box (cyan), UbcH5a (grays) and CHIP U-box (blue white) are shown in cartoon form. **(*C*)** Binding affinity of Avr1d or GmE2 with GmPUB13. ITC binding curve between Avr1d and GmPUB13(252–630) (left). ITC binding curve between GmE2 and GmPUB13(252–630) (right). The ITC experiments were repeated twice independently with similar results.

Avr1d displayed the structural characteristic of a WY motif, adopting a three alpha helix bundle, in which the highly conserved Trp96 and Try118 formed a hydrophobic core. It shared structural similarity with Avr3a11 (3ZR8) (11), another WY motif-containing RxLR effector protein, yielding Cα RMSDs of 1.95 Å, despite low sequence similarity (*SI Appendix,* Fig. S4*B*).

The U-box domain of GmPUB13 consisted of a central α-helix α1, a C-terminal helix α2, a small antiparallel β-sheet (β1 and β2) and two prominent loops (loop1 and loop2) (Fig. 2*A*), resembling the structure of the U-box of the eukaryotic ubiquitin ligase, CHIP (PDB: 2OXQ) (12); it could be superimposed well with the CHIP U-box with a Root-Mean-Square Deviation (RMSD) of 1.66 Å over 72 matching Cα atoms (Fig 2*B*). The hydrophobic groove formed by loop1, loop2 and central α-helix α1 of GmPUB13 U-box contributed to the interaction with Avr1d. Five residues of GmPUB13, P264, I265, L267 (from loop 1), W290 (from helix α1) and P299 (from loop2) formed hydrophobic interactions with five residues of Avr1d, I87, F90, F93 (from helix α2), A117, and I120 (from helix α3). In addition to the hydrophobic interactions, hydrogen bonds and a salt bridge formed between GmPUB13 D259 and Avr1d R123 also stabilized the complex (Fig.3*A*). Intriguingly, the residues in CHIP corresponding to the Avr1d-binding residues of GmPUB13 in CHIP are F218, I216, H241 and P250, and these four residues are necessary for the interaction of CHIP with the E2 subunit, UbcH5a (*SI Appendix,* Fig. S4*C*). Therefore, by similarity to the UbcH5a-CHIP complex, the Avr1d-binding site of GmPUB13 was predicted also be its E2 binding site, and suggesting that Avr1d may compete with E2 for GmPUB13 binding. The isothermal titration calorimetry assay was used to evaluate the binding affinity of GmPUB13-Avr1d and GmPUB13-GmE2. As the result shown in Fig. 2*C*, the binding affinity of GmPUB13-Avr1d (Kd 89 nM) was much stronger than that of GmPUB13-GmE2 (Kd 21 μM). Altogether, these results suggested that Avr1d tightly occupied the E2 binding site of GmPUB13 to compete with E2 for GmPUB13 binding.

**Fig. 3.**
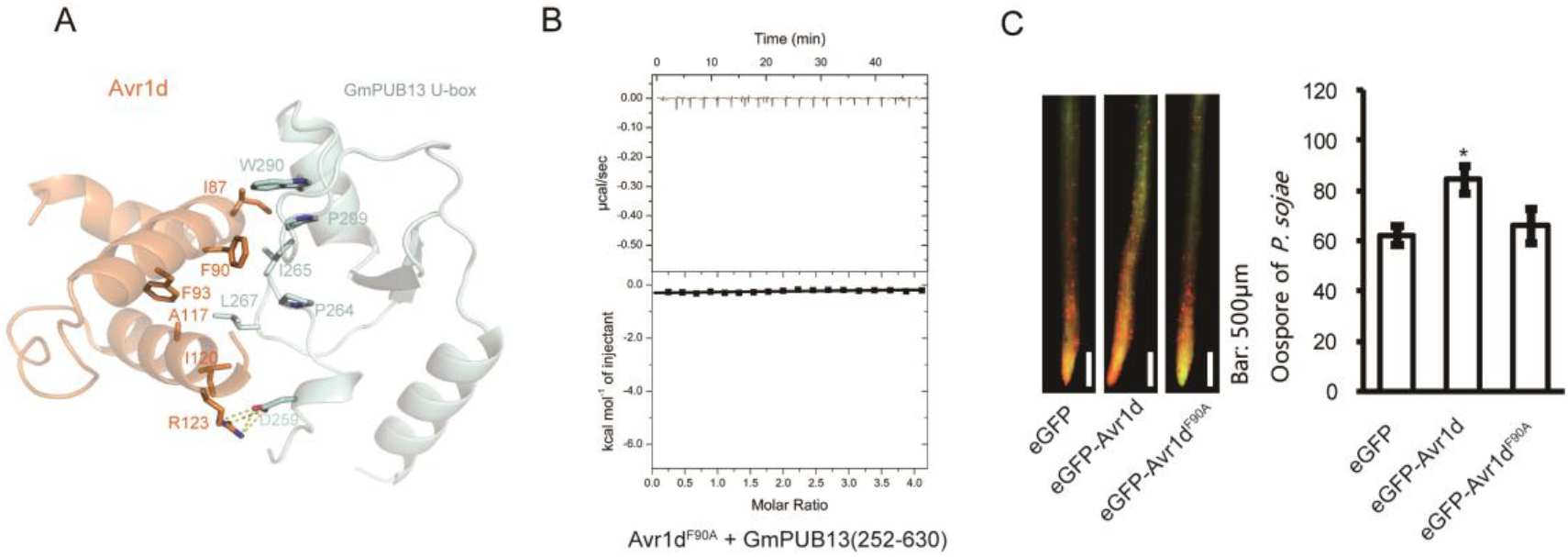
The 90^th^ F of Avr1d is the key residue for the interaction with GmPUB13 and its virulence function. **(*A*)** Interaction residues between Avr1d and GmPUB13. I87, F90, F93, A117 and I120 of Avr1d (orange) contribute to the hydrophobic interaction surface with P264, I265, L267, W290 and P299 of GmPUB13 U-box (cyan). R123 of Avr1d (orange) forms hydrogen bonds and salt bridge with D259 of GmPUB13 U-box (cyan). **(*B*)** ITC binding curve between Avr1d^F90A^ and GmPUB13(252– 630). The ITC experiments were repeated twice independently with similar results. **(*C*)** Overexpression of Avr1d^F90A^ in soybean hairy-roots fails to promote *P. sojae* infection. The fluorescence hairy-roots expressing eGFP, eGFP-Avr1d and eGFP-Avr1d^F90A^ were inoculated with mRFP-labeled *P. sojae* hyphae on the tips (left). Scale bar: 500 μm. The oospore production was observed under fluorescence microscopy at 2 days post inoculation (right). Asterisks indicated t-test p<0.05. These experiments were repeated 3 times with similar results.

### The Phenylalanine 90 of Avr1d is required for interaction with GmPUB13 and virulence function

The crystal structure revealed that hydrophobic interactions played major roles in Avr1d-GmPUB13 complex formation, in which the phenyl ring of Avr1d F90 inserted into the hydrophobic groove formed by P264, I265, W290 and P299 of GmPUB13 (Fig. 3*A*). Substitution of these residues in GmPUB13, P264, I265, W290 and P299, with alanine, compromised Avr1d binding (*SI Appendix,* Fig. S5*A*). On the other side, the mutation F90A in Avr1d likewise disrupted GmPUB13 binding as measured by ITC, gel filtration and yeast two-hybrid assays, indicating that F90 was essential for the interaction with GmPUB13 (Fig. 3*B*, *SI Appendix,* Fig. S5*A*). To evaluate the effect of this mutation on the potential virulence contribution of Avr1d, we overexpressed Avr1d^F90A^ and wild type Avr1d in soybean hairy-roots, then inoculated the roots with mRFP-labeled *P. sojae*. We observed that *P. sojae* oospore production in hairy-roots expressing Avr1d^F90A^ was not significantly different than in hairy-roots expressing eGFP (Fig. 3*C*). Avr1d^F90A^ protein expression in the transformed hairy-roots was confirmed by western blotting with anti-GFP antibodies (*SI Appendix,* Fig. S6). Thus, Avr1d^F90A^ failed to promote *P. sojae* infection. Together, these results demonstrated that F90 of Avr1d was required for the interaction with GmPUB13 and for its potential virulence contribution.

### Avr1d inactivates the ubiquitin ligase activity of GmPUB13 *in vitro*

According to our structure and biochemical assays, Avr1d could compete with E2 for GmPUB13 binding. To determine whether Avr1d affects the ubiquitin ligase activity of GmPUB13, we performed a ubiquitination assay *in vitro*. GST-GmPUB13 or GST-GmPUB13(252-630) proteins were incubated with wheat E1 and Arabidopsis E2 (AtUBC8) together with His-tagged ubiquitin or flag-ubiquitin. The smeared bands detected by ubiquitin antibodies following western blotting indicated that poly-ubiquitin chains of different molecular weights had been formed (Fig. 4*A*). In contrast, no smeared bands were detected in the reaction containing E1, E2, GST or in the reactions containing GmPUB13 but lacking E1 or E2 (Fig. 4*A*), showing GmPUB13 was an active ubiquitin ligase. It has been reported that cysteine 262 and tryptophan 289 of AtPUB13 in the U-box domain are required for its ligase activity (5). GmPUB13 with mutations on these sites, such as GmPUB13^C263A^ or GmPUB13^W290A^ lost ubiquitin ligase activity since no smear bands of poly-ubiquitin were detected in these reactions (Fig. 4*A*), supporting the importance of the U-box domain for enzyme activity. To test whether Avr1d could modulate the enzyme activity of GmPUB13, Avr1d was added into the reaction mixture. The smeared bands were much weaker compared with the control (Fig. 4*B*). Thus, Avr1d indeed inhibited ubiquitin ligase activity of GmPUB13. In line with this, Avr1d decreased the ubiquitin ligase activity of GmPUB13 in a dose-dependent manner since the smeared poly-ubiquitin bands became progressively weaker as the concentration of Avr1d was increased (Fig. 4*B*, *SI Appendix,* Fig. S7*A*). The Avr1d^F90A^ mutant that lost interaction with GmPUB13 could not block the ubiquitin ligase activity of GmPUB13(252-630) (Fig. 4*C*). In addition, increasing the amount of Avr1d^F90A^ protein also failed to block the ubiquitin ligase activity of GmPUB13(252-630) (*SI Appendix,* Fig. S7*B*).

**Fig. 4.**
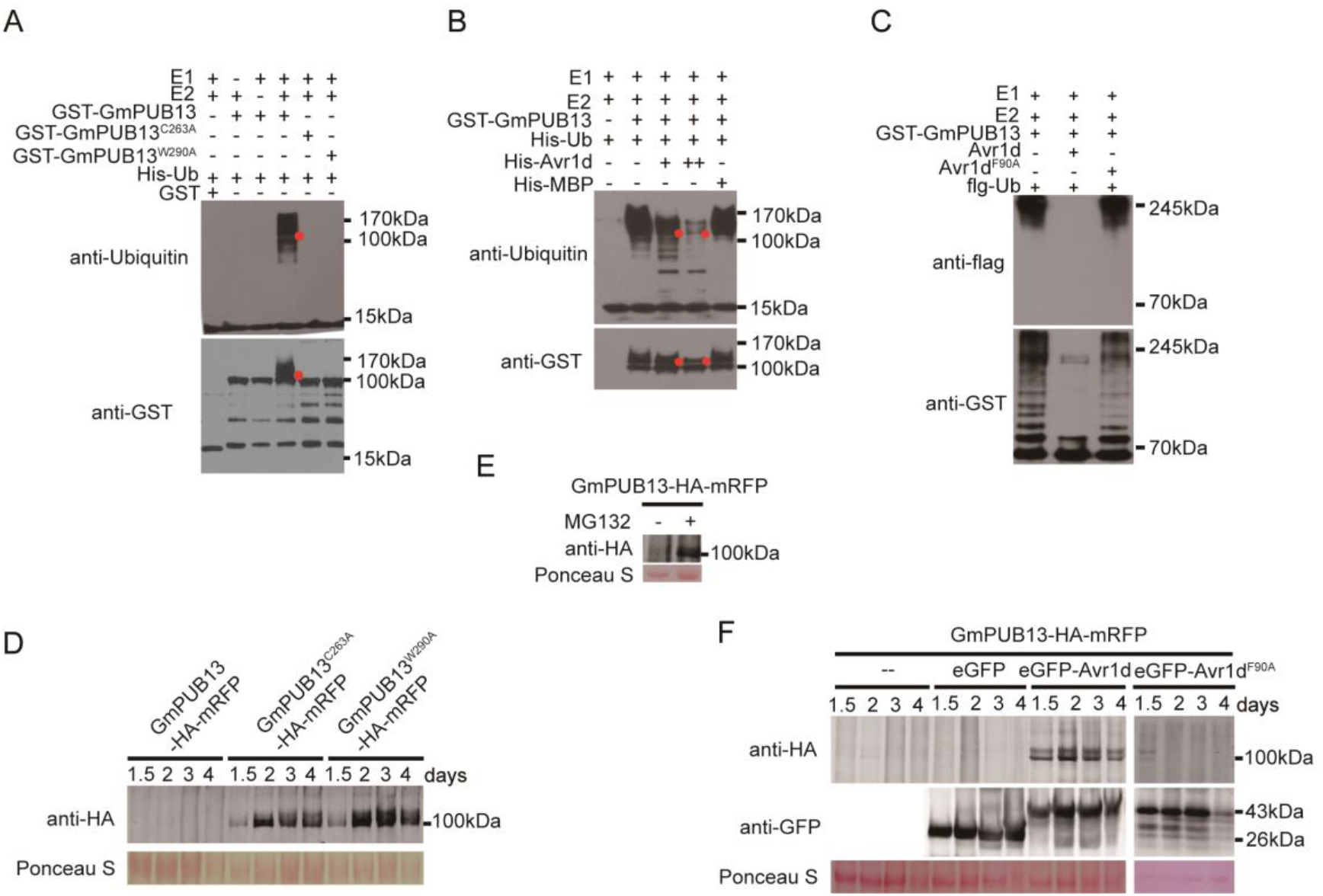
Avr1d inhibits GmPUB13’s ubiquitin ligase activity to promote its accumulation. **(*A*)** Ubiquitination assays of GmPUB13, GmPUB13^C263A^ and GmPUB13^W290A^ *in vitro*. GmPUB13 and mutants were fused with GST-tag and were incubated with or without wheat E1, AtUBC8 as E2 in the buffer with ubiquitin. Reaction mixtures were detected by western blot with anti-ubiquitin and anti-GST. Red dots indicate the protein size of GST-GmPUB13. Molecular mass markers are shown (kDa). **(*B*)** Avr1d inhibits the ubiquitin ligase activity of GmPUB13. His-Avr1d and His-MBP proteins were added into the reaction mixtures with E1, E2 and GST-GmPUB13 as E3. Double plus (++) indicated double amount of His-Avr1d protein. Reaction mixtures were detected by western blot with anti-ubiquitin and anti-GST. Red dots indicate the protein size of GST-GmPUB13. Molecular mass markers are shown (kDa). **(*C*)** Avr1d^F90A^ fails to inhibit the ubiquitin ligase activity of GmPUB13. Avr1d and Avr1d^F90A^ protein were respectively added into the reaction mixture with E1, E2, flag-ubiquitin and GST-GmPUB13(252-630) as E3. Reaction mixtures were detected by western blot with anti-GST and anti-flag. Molecular mass markers are shown (kDa). **(*D*)** Protein accumulation of GmPUB13 and its mutants. GmPUB13-HA-mRFP, GmPUB13^C263A^-HA-mRFP and GmPUB13^W290A^-HA-mRFP were expressed in *N. benthamiana* leaves by agroinfiltration respectively. The total protein were extracted at 1.5, 2, 3 and 4 days post infiltration followed by western blot detection with anti-HA antibody. Ponceau S: ponceau staining indicates the rubisco protein. Molecular mass markers are shown (kDa). **(*E*)** MG132 treatment increases GmPUB13-HA-mRFP protein abundance. The leaves expressing GmPUB13-HA-mRFP were infiltrated with 50 μM MG132 12 hours before harvested and 0.5% DMSO as control. The total protein was detected by western blot with anti-HA antibody. Ponceau S: ponceau staining indicates the rubisco protein. Molecular mass markers are shown (kDa). **(*F*)** Avr1d, but not Avr1d^F90A^ can stabilize GmPUB13. GmPUB13-HA-mRFP was expressed alone or co-expressed with eGFP, eGFP-Avr1d or eGFP-Avr1d^F90A^ in *N. benthamiana* leaves by agroinfiltration. The total proteins were extracted at 1.5, 2, 3 and 4 days post infiltration followed by western blot detection with anti-HA antibody. Ponceau S: ponceau staining indicates the rubisco protein. Molecular mass markers are shown (kDa).

### GmPUB13 self-ubiquitination regulates its abundance

To explore whether GmPUB13 could regulate its protein abundance by self-ubiquitination, we expressed the inactive mutants GmPUB13^C263A^ and GmPUB13^W290A^ in *N. benthamiana* leaves. The protein abundance and fluorescence of GmPUB13^C263A^-HA-mRFP, GmPUB13^W290A^-HA-mRFP and control HA-RFP were much stronger than GmPUB13-HA-mRFP (Fig. 4*D*, *SI Appendix,* Fig. S8*A, B*). The protein abundance of GmPUB13-HA-mRFP was significantly increased when the infiltrated leaves were treated with 50 μM MG132, a 26S proteasome complex inhibitor (Fig. 4*E*). Together, these data suggest that GmPUB13 may regulate its protein abundance by self-ubiquitination and 26S proteasome-mediated degradation.

### GmPUB13 promotes plant susceptibility

To determine the role of GmPUB13 in soybean resistance, we silenced GmPUB13 and GmPUB13L in soybean hairy-roots. The qRT-PCR assay revealed that GmPUB13/GmPUB13L gene expression level decreased around 60% in the silenced hairy-roots with little effect on a homologous gene Glyma10g35220 when compared to the empty vector control (*SI Appendix,* Fig. S9*A, B*). The GmPUB13/GmPUB13L silenced hairy-roots showed less oospore production and *P. sojae* biomass compared to the control hairy-roots when inoculated with eGFP-labeled *P. sojae* (Fig. 5*A*), suggesting that GmPUB13 and GmPUB13L are susceptibility factors that negatively regulate soybean resistance to *P. sojae*. To further verify the conclusion, we overexpressed GmPUB13-HA-mRFP and mutants in *N. benthamiana* and then performed infection assays using *P. capsici*. Compared with the HA-mRFP control, *N. benthamiana* leaves expressing GmPUB13-HA-mRFP showed no difference in lesion area (Fig. 5*B*). However, the leaves expressing GmPUB13^C263A^ or GmPUB13^W290A^ showed much bigger lesions than those expressing GmPUB13 (Fig. 5*C, D*) or HA-mRFP control (Fig. 5*E, F*). These results suggested that the more stable inactivated mutants of GmPUB13 could promote *P. capsici* infection when overexpressed in *N. benthamiana.* These results are consistent with GmPUB13 being a negative regulator of plant immunity, like AtPUB13 (4, 5).

**Fig. 5.**
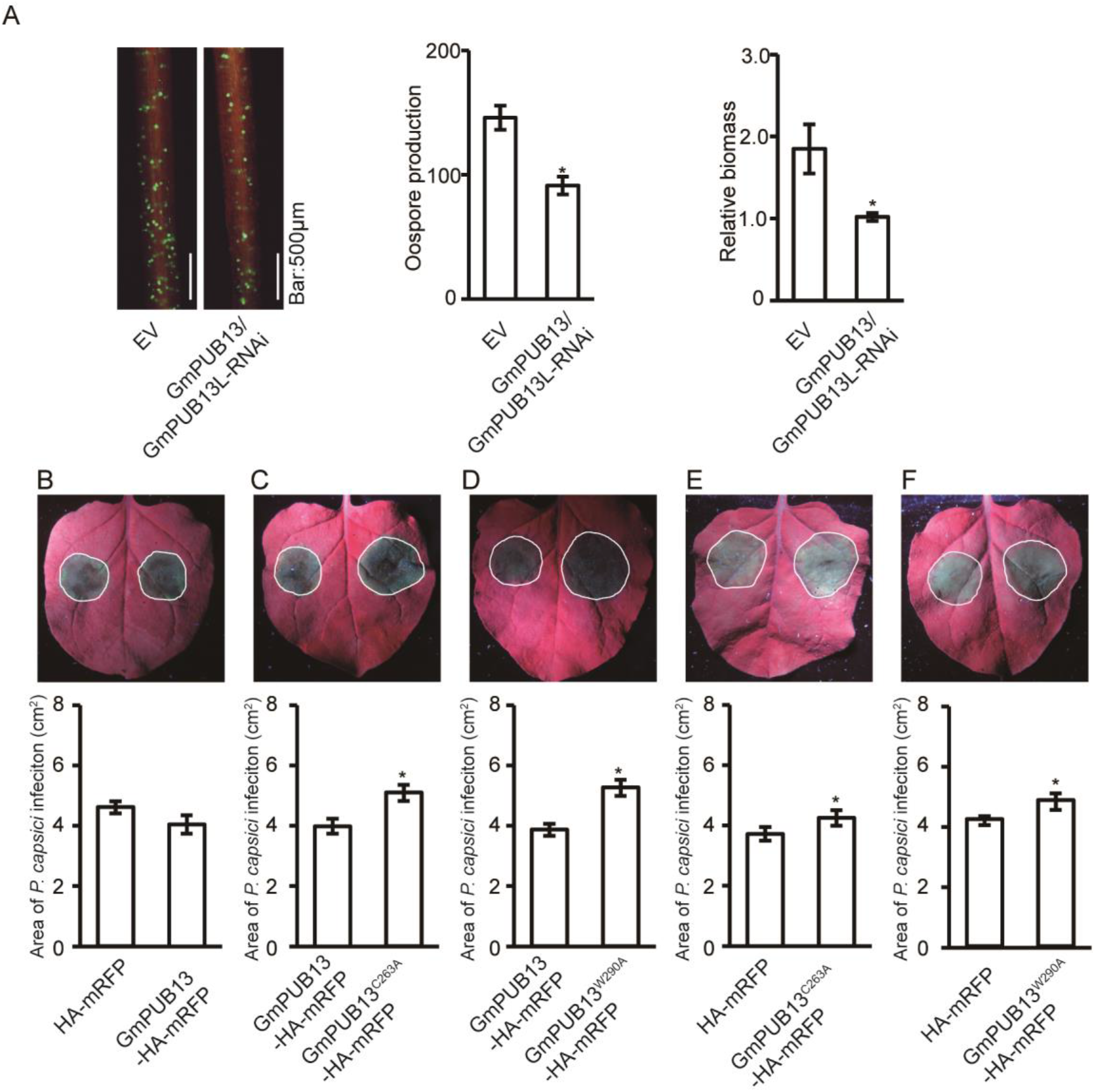
GmPUB13 negatively regulate soybean resistance to *P. sojae*. **(*A*)** Silencing of GmPUB13 and GmPUB13L in soybean hairy-roots by RNAi decreases *P. sojae* infection. GmPUB13 and GmPUB13L were silenced in soybean hairy-roots by RNAi. The tips of red fluorescence hairy-roots were inoculated with eGFP-labeled *P. sojae* hyphae plugs (left). The oospore production was observed under fluorescence microscopy at 2 days post inoculation (middle). The relative *P. sojae* biomass was estimated by DNA-based quantitative PCR (qPCR) (right). Asterisks indicated t-test p < 0.05. These experiments were repeated 3 times with similar results. Bar = 500 μm. **(*B*)** Infection assays on *N. benthamiana* leaves expressing HA-mRFP and GmPUB13-HA-mRFP. HA-mRFP and GmPUB13-HA-mRFP were expressed in each side of the vein of *N. benthamiana* leaves and then inoculated with *P. capsici* plugs. **(*C, D*)** Infection assays on *N. benthamiana* leaves expressing GmPUB13-HA-mRFP and mutants. GmPUB13-HA-mRFP together with GmPUB13^C263A^-HA-mRFP or GmPUB13^W290A^-HA-mRFP were expressed in each side of the vein of *N. benthamiana* leaves respectively and then inoculated with *P. capsici* plugs. **(*E, F*)** Infection assays on *N. benthamiana* leaves expressing HA-mRFP and GmPUB13 mutants. HA-mRFP together with GmPUB13^C263A^-HA-mRFP, or GmPUB13^W290A^-HA-mRFP were expressed in each side of the vein of *N. benthamiana* leaves and then inoculated with *P. capsici* plugs. **(*B - F*)** The dashed circle indicated the lesion of *P. capsici* infection under ultraviolet light. The areas of *P. capsici* infection were shown below. Asterisks indicated t-test p < 0.05. These assays were repeated for 3 times with similar results.

### Avr1d can stabilize GmPUB13 *in planta*

Since GmPUB13 could undergo self-ubiquitination to regulate its protein abundance and Avr1d inhibited the ubiquitination activity of GmPUB13, we tested whether Avr1d could stabilize GmPUB13 *in planta*. When we co-expressed Avr1d and GmPUB13 in *N. benthamiana* leaves, we observed that the fluorescence of GmPUB13-HA-mRFP with eGFP-Avr1d was stronger than those expressing GmPUB13-HA-mRFP alone or GmPUB13-HA-mRFP with eGFP (*SI Appendix,* Fig. S10*A*). We also detected stronger bands of GmPUB13-HA-mRFP in the total proteins of leaves co-expressing GmPUB13-HA-mRFP with eGFP-Avr1d than with eGFP or GmPUB13-HA-mRFP alone for 1.5, 2, 3, 4 days by anti-HA (Fig. 4*F*). Unlike Avr1d, co-expressing Avr1d^F90A^ failed to stabilize GmPUB13 in *N. benthamiana* leaves (Fig. 4*F*, *SI Appendix,* Fig. S10*A*). This is consistent with the previous results in Fig. 4*B, C* suggesting that Avr1d could stabilize GmPUB13.

## Discussion

Plant pathogens secrete large arsenals of effectors to modulate host cell processes and enable a successful infection. Studies on the host targets of such effectors provide molecular clues for understanding the outcome of plant-pathogen interactions. Thus far, multiple host targets have been identified for effectors secreted by oomycete pathogens. However, structural evidence needed to unravel the mechanisms underlying interactions of effectors and host targets is still missing. In this study, we confirmed that *P. sojae* avirulence effector Avr1d physically binds to the soybean U-box type E3 ligase GmPUB13 and we dissected the structural basis of this interaction.

A typical feature of most E3 ligases is the ability to catalyze their own ubiquitination. The biological role of self-ubiquitination was proposed to target the ligase for degradation, which could function as a means of negative feedback of E3 ligases (13, 14). In fact, many E3 ligases, even those that catalyze their own ubiquitination, are targeted by an exogenous ligase, which makes the regulation of E3 ligases rather complicated. In addition, self-ubiquitination of E3 ligases could be involved in non-proteolytic functions (15) and could enhance substrate ubiquitin ligase activity (16). In this study, we found that GmPUB13 was a substrate of its own E3 ubiquitin ligase activity and this self-ubiquitination induced degradation of GmPUB13 by the proteasome.

Ubiquitination contributes crucially to the intricate and precise molecular mechanisms that govern plant immune responses and therefore the ubiquitination system is a hub targeted by multiple pathogen virulence effectors. For example, several effectors possess E3 ubiquitin ligase activity, such as AvrPtoB from *Pseudomonas syringae* (17, 18), XopK from *Xanthomonas oryzae pv. oryzae* (19) that degrade host targets to attenuate plant immunity.

In addition, effectors may hijack the host ubiquitination complex to reprogram the stability of host targets. AvrPiz-t from *Magnaporthe oryzae* interacts with the RING type E3 ubiquitin ligase APIP6, a positive regulator of rice immunity, to promote the degradation of APIP6 (20). The *P. infestans* RxLR effector Pi02860 interacts with a potato susceptibility factor StNRL1, a putative substrate adaptor of a CULLIN3-associated ubiquitin E3 ligase complex, to degrade a guanine nucleotide exchange factor called SWAP70, which is essential for potato immunity (21, 22). In addition, *P. infestans* effector AVR3a interacted with a U-box type ubiquitin E3 ligase CMPG1 and manipulated plant immunity by stabilizing potato CMPG1 (23). Nevertheless, how AVR3a mediates the stability of CMPG1 is thus far unclear.

In this study, we found that the *P. sojae* avirulence effector Avr1d physically binds to the soybean U-box type E3 ligase GmPUB13 and could stabilize GmPUB13 by suppressing its ubiquitin ligase activity. We deciphered the molecular mechanism underlying inhibition of GmPUB13 by Avr1d by determining the molecular structure of the Avr1d-GmPUB13 complex, combined with biochemical assays. Based on the crystal structure, we successfully identified the interface required for interaction between GmPUB13 and Avr1d and showed that the residues at the interface were required for Avr1d to bind to GmPUB13 and inhibit its ligase activity. Since GmPUB13 functions as a susceptibility factor, stabilization of GmPUB13 by Avr1d-mediated inhibition of its ligase activity is the key to the contribution of Avr1d to *P. sojae* virulence.

## Acknowledgements

We thank the BL17U1 and BL19U1 of Shanghai Synchrotron Radiation Facility for support during crystallographic data collection. We thank W.Ma (University of California Riverside) for discussions and constructive suggestions. Research in the W.X. laboratory is supported by the Chinese Thousand Talents Plan and the Chinese Academy of Sciences. This research was supported by grants to YC.Wang from the China National Funds for Innovative Research Groups (31721004), the Chinese Modern Agricultural Industry Technology System (CARS-004-PS14), the key program of the National Natural Science Foundation of China (31430073) and the Fundamental Research Funds for the Central Universities (KYXK202010). B.M.T. was supported by Oregon State University.

## Materials and methods

### *Phytophthora*, bacteria and plant growth condition

*Phytophthora sojae* and *P. capsici* were grown in 10% (V/V) V8 medium with 1.5% agar or without agar in dark at 25°C. *Escherichia coli* JM109 for vector construction were grown on LB medium with agar or sharking 220 rpm in dark without agar, 37°C. *Agrobacterium tumefaciens* and *A. rhizogenes* were grown on LB medium with 1.5% agar or with sharking 220 rpm in dark without agar containing 50 μg/L Rifampicin or streptomycin respectively, at 30°C. *Nicotiana benthamiana* were grown on vermiculite from seeds in climate chamber under long day conditions (14 hours light and 10 hours dark) at 24°C. *Glycine max* etiolated seedlings were grown on vermiculite medium from seeds in dark, at 25°C for 3 - 4 days in climate chamber.

### *P. sojae* transformation, zoospore producing and infection assay

Avr1d-mRFP driven by the Ham34 promoter (pTORmRFP4) was transformed into *P. sojae* strain P6497 using PEG-mediated protoplast transformation system (24). Transformants were selected on 10% (V/V) V8 medium containing 50 μg/L geneticin (G418) and further confirmed by red fluorescence selection under fluorescence microscopy and western blot detection of Avr1d-mRFP protein by anti-mRFP antibody. *P. sojae* zoospores were produced by washing the 3 - 5 day old *P. sojae* hyphae grown in 10% V8 liquid medium with sterilized tap water for 3 times and then incubated at dark (25°C) to stimulate sporulation and zoospore releasing. The droplets containing about 100 zoospores were then inoculated to the hypocotyl of soybean etiolated seedlings. The inoculated samples were kept in wet and dark conditions at 25°C. For 5 - 10 hours infection, the epidermal cells of soybean hypocotyl were observed under confocal microscopy.

### Hairy-roots transformation and infection assay

Soybean hairy-root transformation was performed by using *A. rhizogenes* K599 - mediated T-DNA transformation (25). To express Avr1d and its mutant in soybean hairy-root, Avr1d and its mutant were fused with N-terminus eGFP and cloned into the T-DNA region of pBinGFP2 plasmid with CaMV 35S promoter. The green fluorescence hairy-roots selected under fluorescence microscopy were regarded as transformed hairy-root. The protein expression of the fluorescence hairy-roots were detected by western-blot with anti-GFP antibody.

For silencing of *GmPUB13* and *GmPUB13L* in soybean hairy-roots, a 100 bp cDNA sequence that specifically matching to GmPUB13 and GmPUB13L (blasted by the VIGS tool in http://solgenomics.net/) was constructed with forward and reverse sequence linked by chalcone synthase intron from *Petunia hybrida* into pFGC5941mCherry plasmid with CaMV 35S promoter. The T-DNA region contains another open reading frame for mCherry expression driven by mannopine synthase promoter. The red fluorescence hairy-roots under fluorescence microscopy were regarded as silenced hairy-roots. The silencing efficiency was detected by real-time relative quantitated PCR (qPCR) by relative to the gene expression of the soybean CYP2 (8).

To test the susceptibility, the fluorescence hairy-roots were inoculated with eGFP or mRFP labeled *P. sojae* hyphae at the tips and incubated in wet condition and dark, 25°C (26). For 2 days infection, the roots were observed under fluorescence microscopy. The relative biomass of *P. sojae* grown in transformed hairy-roots were detected by real-time quantitative PCR and indicated by the ratio of *P. sojae* actin gene and soybean *CYP2* gene.

### Yeast two-hybrid

The PEG-mediated transformation protocol described in Clontech Yeast Protocol Handbook was used to transform plasmids into yeast AH109 and Matchmaker GAL4 Two-Hybrid System to screen for targets or check the protein - protein interaction. To screen host targets of Avr1d, Avr1d without signal peptide was constructed into vector pGBKT7 as the bait. cDNA library derived from soybean hypocotyl and roots RNA were constructed to pGADT7 plasmid fused with GAL4 activation domain (AD) by Clontech company. A total of 6 × 10^6^ clones was screened. The yeast clones were selected on SD/-His/-Leu/-Trp medium for medium stringency interaction or on SD/-Ade/-His/-Leu/-Trp/X-α-Gal medium for high stringency interaction. The AD plasmids of the selected clones were extracted by Solarbio Plasmid Extraction Mini Kit (Cat: D1100) and were subsequently transformed into *E. coli* JM109 before sending to sequence. The obtained sequences were used to search in phytozome (https://phytozome.jgi.doe.gov/pz/portal.html) database for candidate target genes. To confirm the interaction of Avr1d and full length GmPUB13, Avr1d was constructed into pGBKT7 plasmid, while full length GmPUB13 and GmPUB13L were constructed into pGADT7 plasmid from soybean cDNA. The plasmids were co-transformed into yeasts and then tested on SD/-Leu/-Trp medium and SD/-Ade/-His/-Leu/-Trp/X-α-Gal medium.

### GST-Pull down and Co-IP assay

For GST-Pull down, Avr1d was cloned into pGEX 4T-2 with a C-terminus GST tag, while GmPUB13 was cloned into pET28a fused with His-tag. Both Avr1d and GmPUB13 were expressed in *Escherichia coli* BL21, induced by 0.2 mM IPTG. The bacteria lysate containing GST-Avr1d was incubated with 10 μl glutathione sepharose 4B beads (Cat: 45-000-139, GE Healthcare) for 3 hours and purified by washing with 1×PBS for 5 times. Then the beads were incubated with bacteria lysate containing His-GmPUB13 for another 3 hours and purified by washing with PBS for 5 times. The proteins eluted from the beads were determined by western blot with anti-GST and anti-His. For Co-IP, Avr1d without signal peptide was cloned into pBinGFP2 plasmid fused with eGFP at its C-terminal and GmPUB13 was cloned into pFGC5941HAmRFP fused with 3 × HA-mRFP at its C-terminal. Avr1d and GmPUB13 were co-expressed in *N. benthamiana* leaves by *A. tumefaciens*-mediated transformation. The total protein of *N. benthamiana* leaves was extracted in IP buffer (50 mM Tris-HCl pH=7.5, 150 ml NaCl, 10% glycerol, 0.1% TRITON X-100, 1% cocktail, 1 mM PMSF) and then was incubated for 4 hours with 10 μl agarose beads conjugated anti-HA mouse monoclonal antibody (Cat: AT0079, CMC tag). After washing with 1×PBS buffer for 5 times, the agarose beads with proteins were boiled in 1×SDS-loading buffer for 10 min. Then the protein samples were determined by western blot with anti-HA (Abmart) and anti-GFP (CMC tag).

### Relative quantification of *P. sojae* biomass

The genome DNA of infected soybean hairy-roots was extracted using New Plant Genome Extraction Kit (Cat: DP305, Tiangen biotech) and used for qPCR with ChamQ SYBR qPCR Master Mix (Cat: Q311, Vazyme). The *P. sojae* biomass was calculated by using *P. sojae PsActin* gene relative to soybean *GmCYP2* gene. Primer sequences are listed in supplementary table 2.

### Transient expression in *N. benthamiana*

Avr1d, GmPUB13 and related mutants sequences were amplified by Phanta Max Super-Fidelity DNA Polymerase (Cat: P505, Vazyme) from cDNA of *P. sojae* or *G. max*, respectively. The fragments were then cloned into vector pBinGFP2 or pFGC5941HAmRFP using ClonExpress II One Step Cloning Kit (Cat: C112, Vazyme). *A. tumefaciens* strains with different plasmids were infiltrated into *N. benthamiana* leaves. The leaves were harvested 1.5, 2, 3, 4 days post infiltration. Total proteins were extracted in IP buffer (50 mM Tris-HCl pH=7.5, 150 ml NaCl, 10% glycerol, 0.1 - 1% Triton X-100, 1% cocktail, 1 mM PMSF). The proteins of different samples were determined by western blot with anti-HA or anti-HA (Abmart) and anti-GFP (CMC tag).

### MG132 treatment

MG132, a 26S proteasome inhibitor, was dissolved in DMSO with a concentration of 10 mM. MG132 diluted in 10 mM MgCl_2_ buffer with a final concentration of 50 μM was infiltrated into *N. benthamiana* leaves 12 hours before harvested. 5% DMSO diluted in 10 mM MgCl_2_ buffer was infiltrated as control.

### Inoculation with *P. capsici* on *N. benthamiana* leaves

*A. tumefaciens* strain pairs for expressing HA-mRFP and GmPUB13, HA-mRFP and GmPUB13 mutants or GmPUB13 and mutants were infiltrated into the two sides of *N. benthamiana* leaves (>18 leaves), respectively. Then 36 hours post infiltration, infiltrated leaves were inoculated with mycelia plugs of fresh *P. capsici* (plug diameter was about 5mm) and kept in dark with high humidity for 2 days. The disease symptoms were visualized under ultraviolet light.

### Ubiquitination assay *in vitro*

Ubiquitination assay *in vitro* was carried out following the protocol described by Zhao (27). GmPUB13 or GmPUB13(252-630) and other mutants fused with GST-tag were expressed in *E. coli* BL21 and were purified by immunoprecipitation with glutathione sepharose 4B beads (Cat: 45-000-139, GE Healthcare). Each reaction with 30 μl final volume contained 50 ng Wheat E1 protein, 200 - 500 ng AtUBC8 protein as E2 conjugating enzyme, 5 mg ubiquitin together with reaction buffer (50 mM Tris-HCl pH 7.4, 10 mM MgCl_2_, 5 mM dithiothreitol, 5 mM ATP, 10% glycerol) in the tubes containing beads binding with GmPUB13, mutants or GST-tag proteins. The reactions were incubated in thermomixer for 3 hours, 30°C, with shaking, and stopped by adding 1 × SDS-PAGE loading buffer and incubated for another 5 min at 100°C. Avr1d and Avr1d^F90A^ aliquoting (5 μl) for each reaction were analyzed by electrophoresis on 12% SDS-PAGE gels. The incubated mixtures were detected by western blot using anti-ubiquitin (Abcam), anti-GST (Abmart) and anti-Flag (Abmart).

### Protein Expression and Purification

The GmPUB13 U-box and Avr1d were both cloned into the modified pET32a vector (Novagen) after adding a cleavage site for the tobacco etch virus (TEV) protease to the 5’ and 3’ ends of the Avr1d and GmPUB13 gene through PCR amplification. For His-GmPUB13 U-box and Avr1d protein co-expression, the construct was transformed into *E. coli* BL21(DE3), and cells were grown at 37°C to OD_600_ = 0.6 - 0.8 and induced with 0.1 mM isopropyl b-D-1-thiogalactopyranoside for 8-10 hours at 16°C. The cells were harvested by centrifugation for 15 min at 4000 g at 4°C. Cells were resuspended in Ni-lysis buffer (50 mM Tris [pH 8.0], 200 mM NaCl, 20 mM imidazole) and lysed with a homogenizer. The lysate was centrifuged for 1 hour at 17000g (4°C), and the supernatant was passed over a Ni-affinity column (GE Healthcare). His-GmPUB13 U-box and Avr1d were recovered by gradient elution with Ni-elution buffer (50 mM Tris [pH 8.0], 200 mM NaCl, 200 mM imidazole). Fractions containing His-GmPUB13 U-box and Avr1d were verified by SDS-PAGE and pooled for tag cleavage overnight at 4°C with 1:20 tobacco etch virus (TEV) protease. Untagged His-GmPUB13 U-box and Avr1d were loaded on a superdex 200 gel-filtration column (GE Healthcare) and eluted in superdex buffer (20 mM Tris [pH 8.0], 300 mM NaCl, 1 mM TCEP) using the AKTA Explorer FPLC system (GE Healthcare). The protein was concentrated to 7.5 mg/ml.

### Crystallization, data collection and structure determination

Crystallization was conducted using the sitting-drop vapor diffusion method at 4°C. Avr1d-GmPUB13 U-box yielded crystals with good diffraction quality in 12% w/v Polyethylene glycol 20,000, 0.1 M BICINE pH 8.5 and 3% w/v Dextran sulfate sodium salt. Before data collection, all crystals were soaked in the reservoir solution supplemented with 25% glycerol and flash-cooled in liquid nitrogen. The diffraction data were collected at beamlines BL17U1 and BL19U1 at the Shanghai Synchrotron Radiation Facility and were processed using the XDS program. A summary of the statistical methods used for data collection and analysis is provided in Supplementary Table1. Phases were obtained experimentally with data from selenomethionine-substituted Avr1d-GmPUB13 U-box. The PHENIX software suite was used for initial model building. The final model was built by performing iterative manual model building with Coot and maximum likelihood refinement in PHENIX. Images and structural alignments were generated by using PyMOL.

### Size exclusion chromatography (SEC) assay

Purified GmPUB13(252-630) (roughly 40 μM) and Avr1d (roughly 40 μM) proteins were incubated at 4°C for one hour in the buffer containing 20 mM Tris [pH 8.0], 300 mM NaCl, 1 mM TCEP. GmPUB13(252-630), Avr1d, GmPUB13(252-630) and Avr1d, three samples were then injected onto a Superdex 200 gel-filtration column (GE) for analysis in line at a flow rate of 0.4 ml/min. The fractions (0.5 ml/fraction) were analyzed by SDS-PAGE electrophoresis and visualized by coomassie brilliant blue staining. Mutant samples such as Avr1d^F90A^ were used the same method as described above.

### Isothermal titration calorimetry

ITC binding curves were measured by using a Microcal PEAQ-ITC instrument (Malvern). Purified proteins were transferred to buffer containing 20mM HEPES pH 7.5, 100mM NaCl, and 2mM β-mercaptoethanol by 5ml desalting column (GE). Titrations were performed at 20°C. Titration of GmPUB13(252-630) in the cell was performed by sequential addition of Avr1d, GmE2 and Avr1d^F90A^ separately. Data were analyzed using Origin 7.0.

## Supplementary information

**Figure S1.**
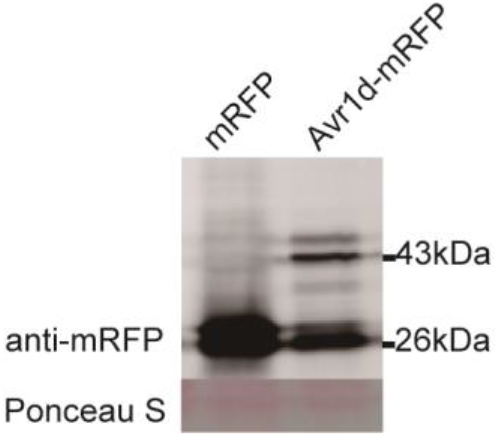
Avr1d-mRFP was overexpressed in *P. sojae*. Avr1d with native signal peptide fused with mRFP at the C-terminus was overexpressed in *P. sojae*. The total proteins were detected by western blot with anti-mRFP. Ponceau S: ponceau staining. Molecular mass markers are shown (kDa).

**Figure S2.**
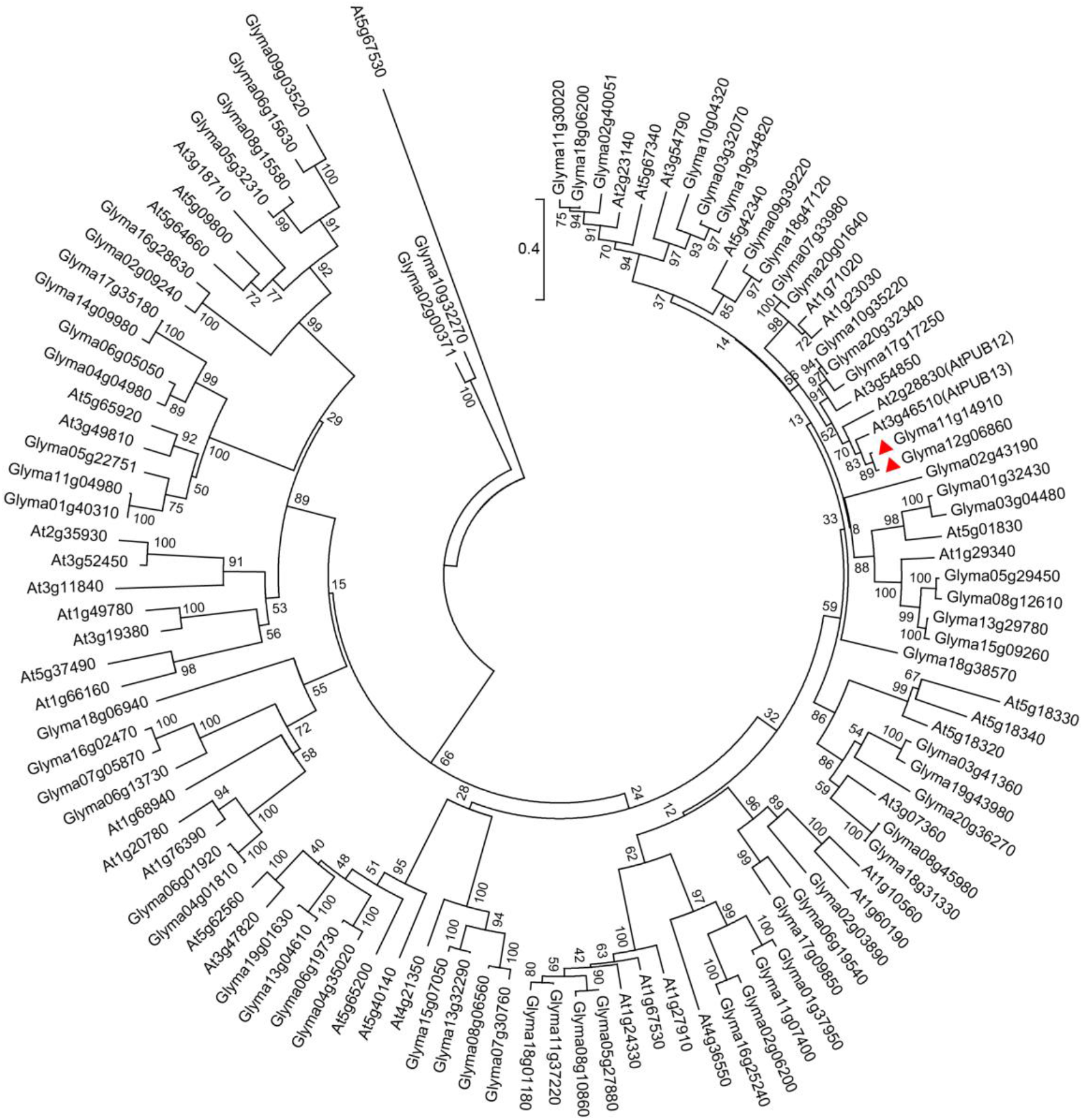
The phylogeny of ARM-repeats type U-box proteins in soybean and *A. thaliana*. A phylogenetic tree of ARM-repeats type U-box proteins in soybean and *A. thaliana* aligned with Clustal W was built by using the method of neighbor-joining in MEGA6 with 1,000 bootstrap replicates. Glyma12g06860 and Glyma11g14910 are marked by red triangles. The branch lengths in the same units were regarded as the evolutionary distances and used to infer the phylogenetic tree. The number on the node shows the confidence level.

**Figure S3.**
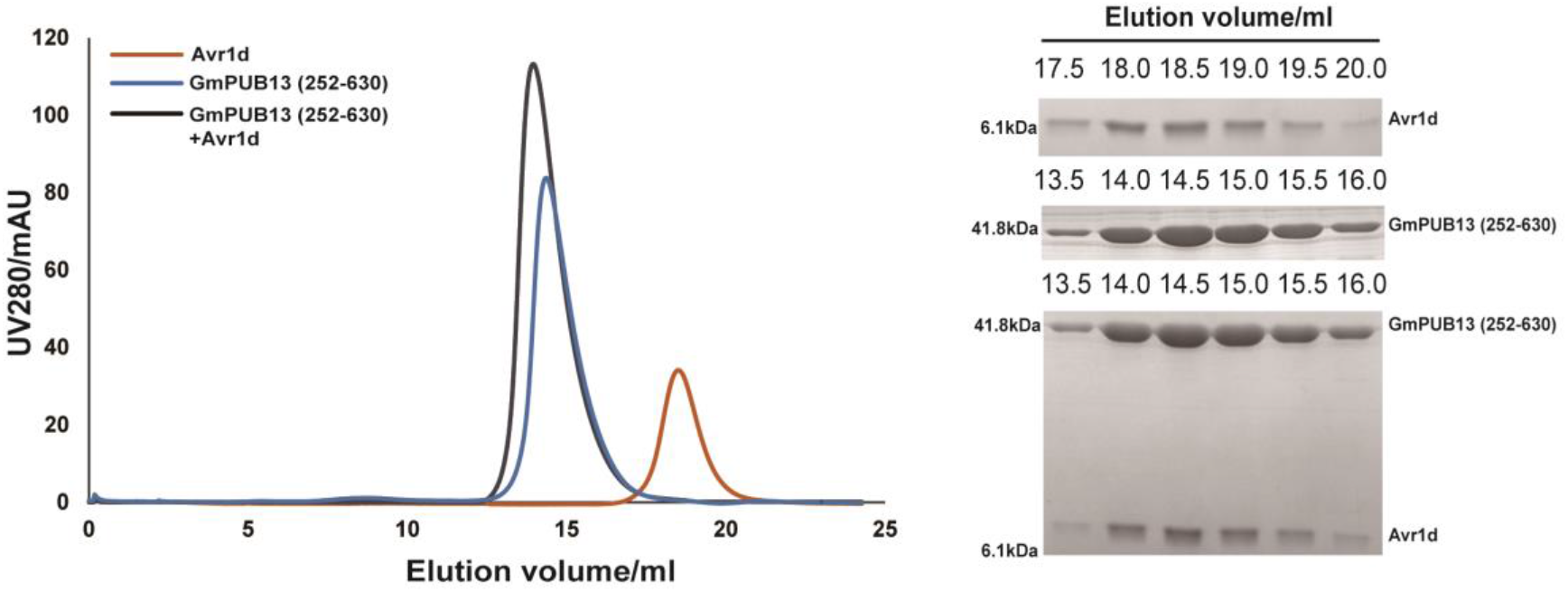
Gel filtration assays demonstrate the interaction between Avr1d and GmPUB13. Gel filtration traces show the retention volume (left) and SDS-PAGE gels (right) of relevant elution fractions of Avr1d, GmPUB13(252-630) and 1:1 mixtures of both proteins. Molecular mass markers are shown (kDa).

**Figure S4.**
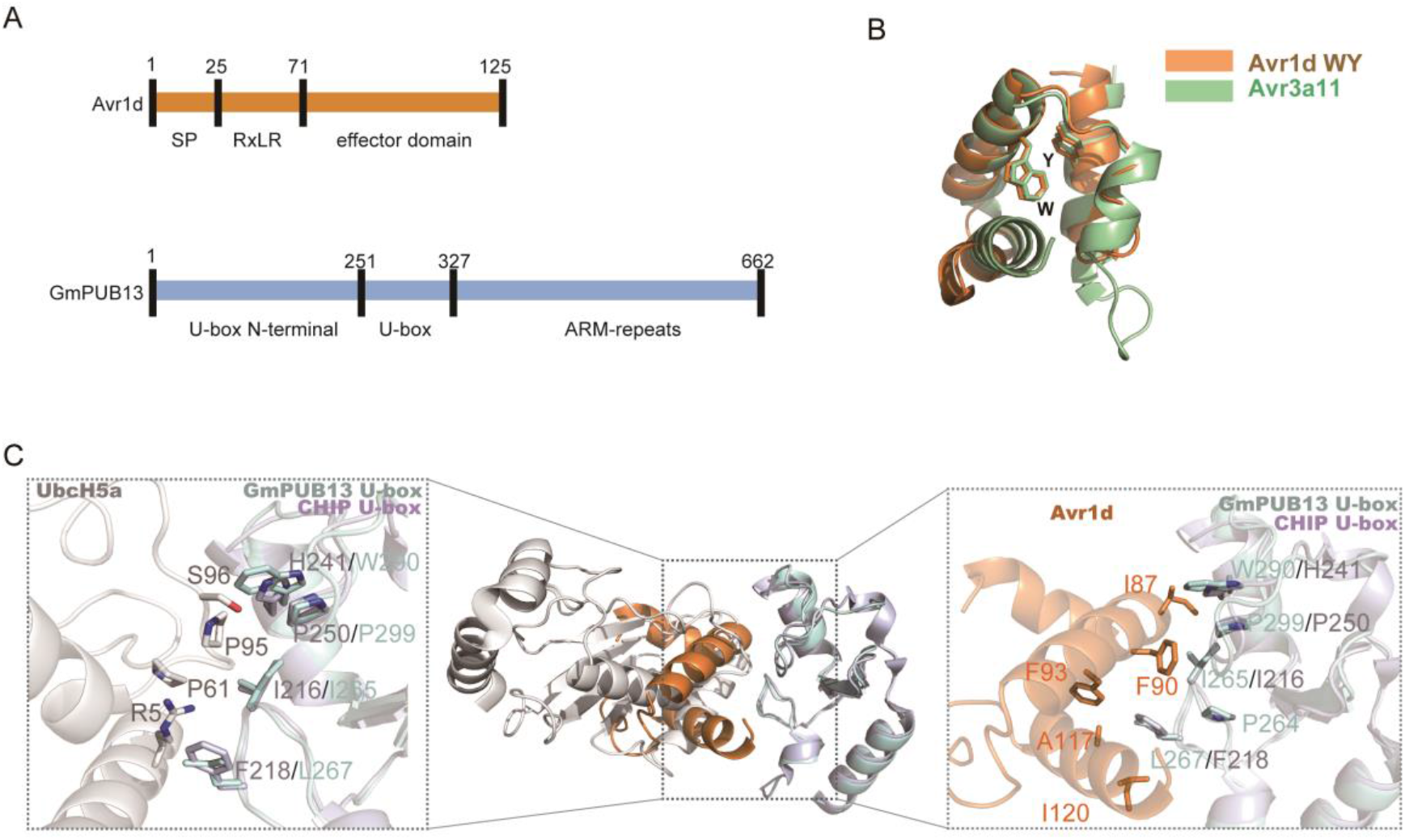
Avr1d occupies the E2 binding site of GmPUB13. **(*A*)** Schematic view of Avr1d and GmPUB13. Avr1d contains a signal peptide, an RxLR motif and an effector domain. GmPUB13 contains a U-box N-terminal domain, a U-box domain, and an ARM-repeats domain. The numbers indicate the sequence number of amino acid residue. **(*B*)** Structure alignment of Avr1d and Avr3a11. Avr1d contains a WY motif. The WY motif adopts a similar structure as Avr3a11, with Cα RMSD values of 1.95 Å. **(*C*)** Superimposition of the Avr1d-GmPUB13 U-box and UbcH5a-CHIP U-box (Avr1d in orange, GmPUB13 U-box in cyan, UbcH5a in gray and CHIP U-box in blue white). The conformations of GmPUB13 U-box and CHIP U-box backbone and interface residues are almost identical between the structures. P264, I265, W290 and P299 of GmPUB13 U-box (cyan), which are necessary for the interaction with Avr1d (orange) correspond to F218, I216, H241 and P250 residues of CHIP U-box (blue white), which are necessary for the interaction with UbcH5a (grays).

**Figure S5.**
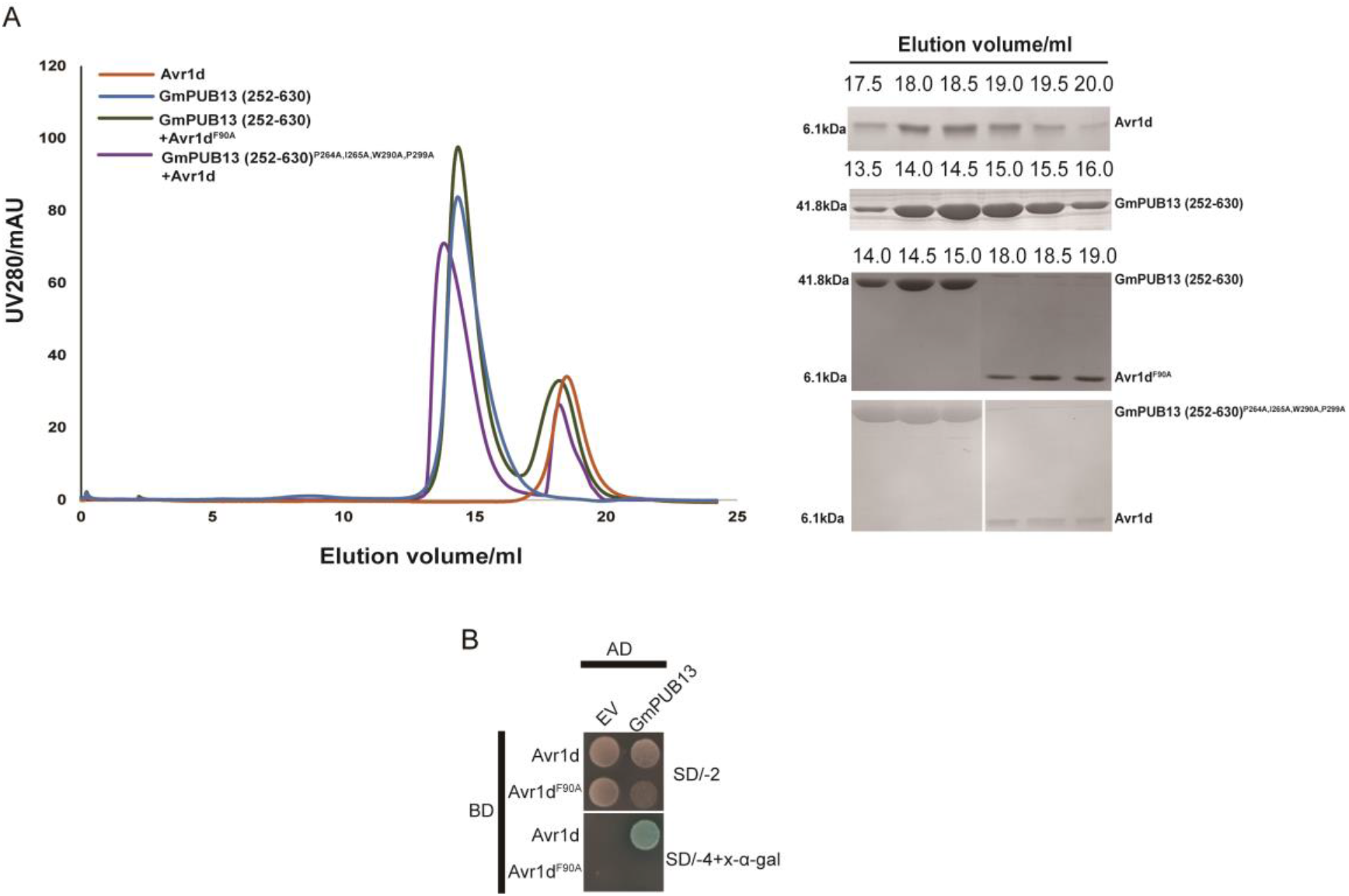
Mutations of Avr1d or GmPUB13 compromise their interaction. **(*A*)** Gel filtration traces show the retention volume (left) and SDS-PAGE gels (right) of relevant elution fractions of Avr1d, GmPUB13(252-630), Avr1d^F90A^:GmPUB13(252-630) and Avr1d:GmPUB13(252-630)^P264A,I265A,W290A,P299A^. Molecular mass markers are shown (kDa). **(*B*)** Yeast two-hybrid assay of Avr1d and Avr1d^F90A^ with GmPUB13. The 90^th^ Phenylalanine residue of Avr1d was mutated to Alanine. pGBKT7::Avr1d or pGBKT7::Avr1d^F90A^ were co-transformed into yeasts with pGADT7::GmPUB13. The interactions were tested by assessing the growth of the yeast on SD/-2 medium (SD/-Leu/-Trp) and SD/-4 + X-α-gal medium (SD/-Ade/-His/-Leu/-Trp + X-α-gal). EV: empty vector.

**Figure S6.**
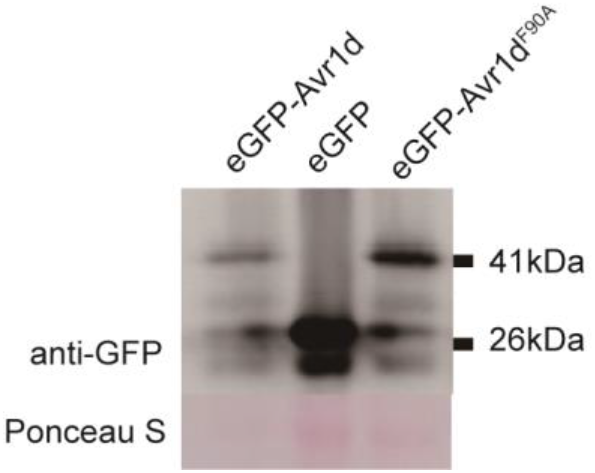
Avr1d and Avr1d^F90A^ were expressed in soybean hairy-roots. eGFP-Avr1d, eGFP and eGFP-Avr1d^F90A^ were expressed in soybean hairy-roots by *A. rhizogenes*-mediated transformation. The total proteins of eGFP-Avr1d, eGFP and eGFP-Avr1d^F90A^ hairy-roots were extracted and detected by western blot with anti-GFP antibody. Ponceau S: ponceau staining. Molecular mass markers are shown (kDa).

**Figure S7.**
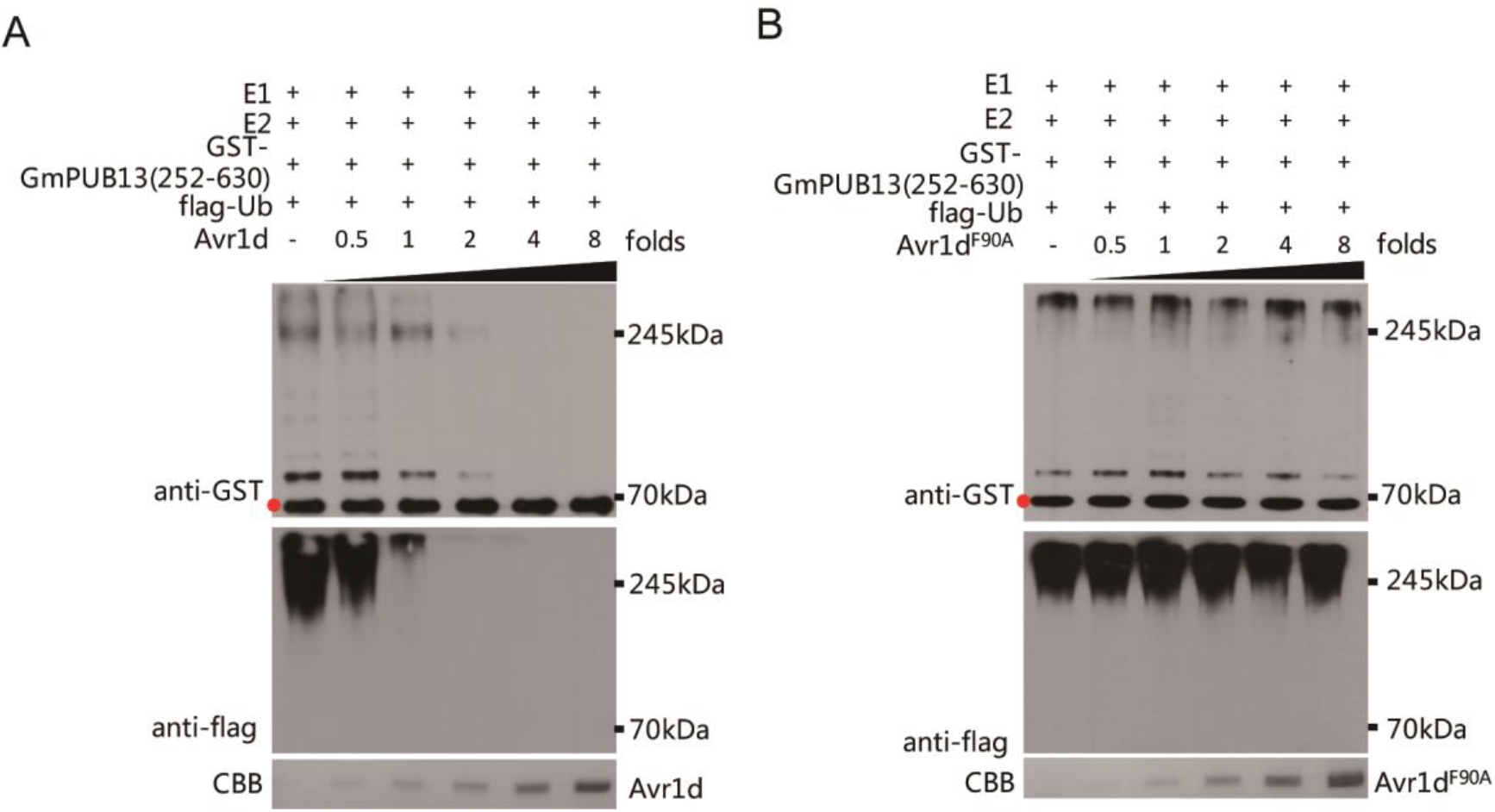
Avr1d, but not Avr1d^F90A^ can block the ubiquitin ligase activity of GmPUB13(252-630) in a dosage manner. **(*A, B*)** The effect of Avr1d and Avr1d^F90A^ to the ubiquitin ligase activity of GmPUB13(252-630). Avr1d or Avr1d^F90A^ protein of 0, 0.5, 1, 2, 4, 8 folds of the amount of GST-GmPUB13(252-630) were added into the reaction mixture with E1, E2, flag-ubiquitin and GST-GmPUB13(252-630) as E3. Reaction mixtures were detected by western blot with anti-flag and anti-GST. Red dots indicated the protein size of GST-GmPUB13(252-630). CBB: Coomassie brilliant blue staining. Molecular mass markers are shown (kDa).

**Figure S8.**
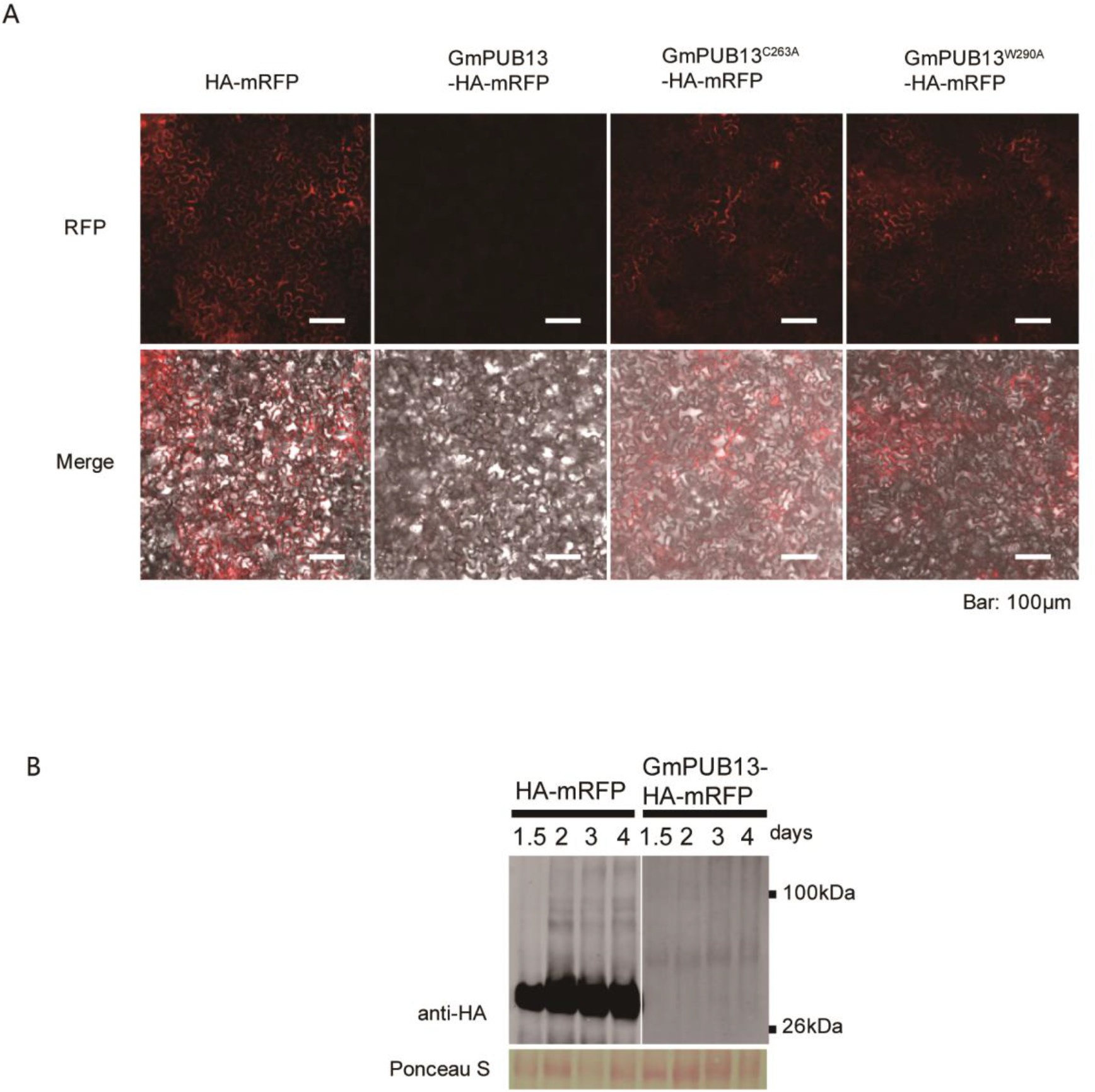
GmPUB13 inactivated mutants are more stable when expressed in *N. benthamiana.* **(*A*)** Fluorescence microscopy view of mRFP, GmPUB13-HA-mRFP and mutants. mRFP, GmPUB13-HA-mRFP, GmPUB13^C263A^-HA-mRFP and GmPUB13^W290A^-HA-mRFP were expressed in *N. benthamiana* leaves by agroinfiltration. Infiltrated leaves were observed under fluorescence microscopy 2 days post infiltration. Scale bar = 100 μm. **(*B*)** GmPUB13 protein was unstable when expressed in *N. benthamiana*. HA-mRFP and GmPUB13-HA-mRFP were expressed in *N. benthamiana* leaves by agroinfiltration. Total protein was extracted 1.5, 2, 3 and 4 days post infiltration followed by western blot detection with anti-HA. Ponceau S: ponceau staining indicates the rubisco protein. Molecular mass markers are shown (kDa).

**Figure S9.**
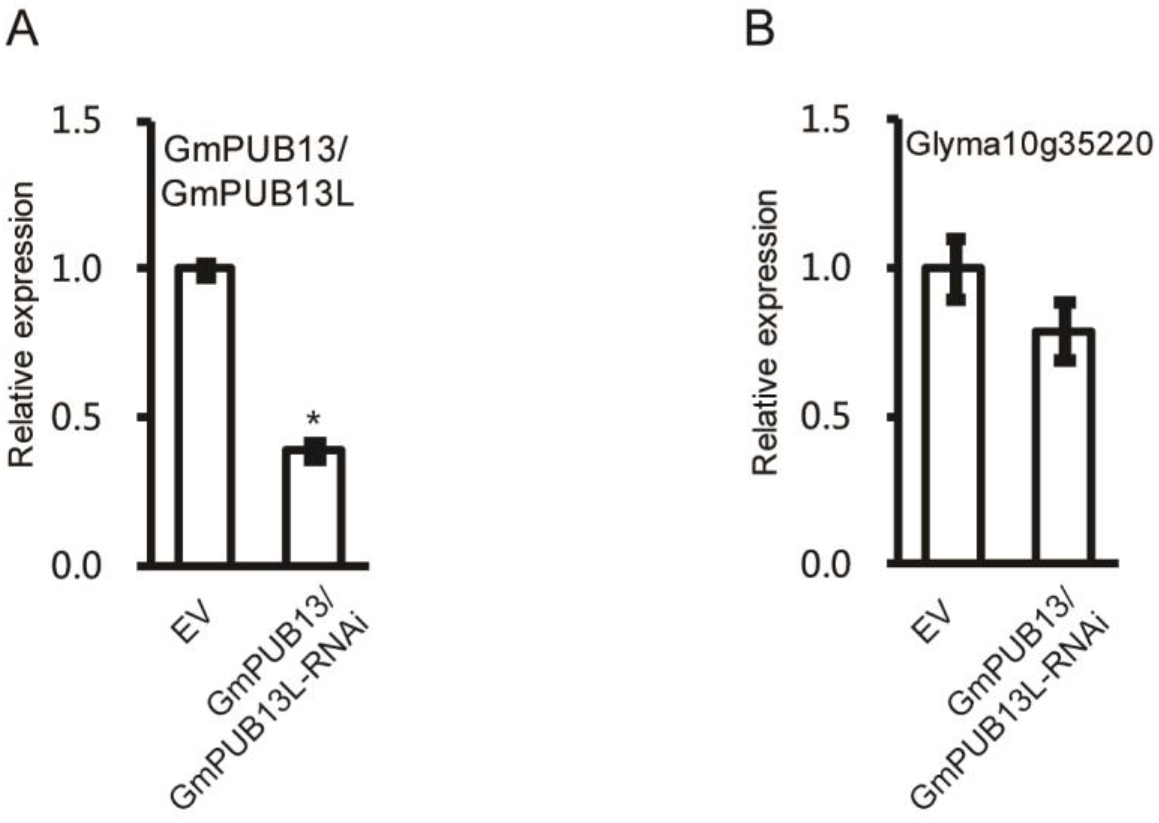
Silencing of GmPUB13 and PUB13L in soybean hairy-roots by RNAi. **(*A*)** GmPUB13 and GmPUB13L were silenced in soybean hairy-roots by RNAi. The silencing efficiency was detected by quantitative PCR (qPCR) and indicated by the ratio of GmPUB13 and GmPUB13L expression versus the soybean CYP2 gene expression. Asterisks indicated t-test p < 0.05. **(*B*)** The effect to the transcription of Glyma10g35220 in GmPUB13/GmPUB13L-RNAi roots was detected by quantitative PCR (qPCR) and indicated by the ratio of Glyma10g35220 expression versus the soybean CYP2 gene expression.

**Figure S10.**
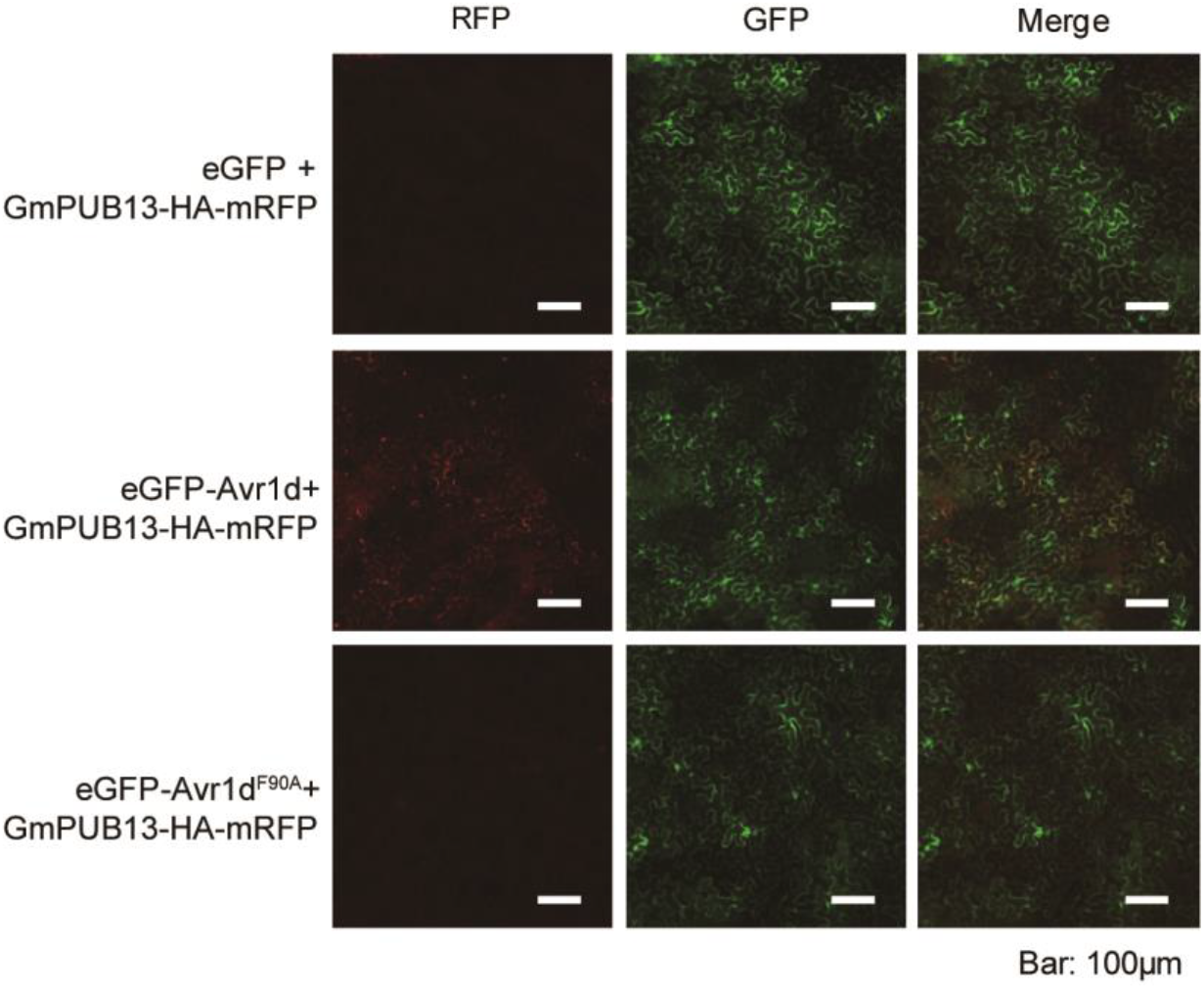
Avr1d, but not Avr1d^F90A^ can stabilize GmPUB13. Fluorescence microscopy views of GmPUB13-HA-mRFP co-expressing with eGFP, eGFP-Avr1d or eGFP-Avr1d^F90A^. GmPUB13-HA-mRFP was co-expressed with eGFP, eGFP-Avr1d and eGFP-Avr1d^F90A^ in *N. benthamiana* leaves by agroinfiltration. Infiltrated leaves were observed under fluorescence microscopy 2 days post infiltration. Scale bar = 100 μm.

**Table S1.**
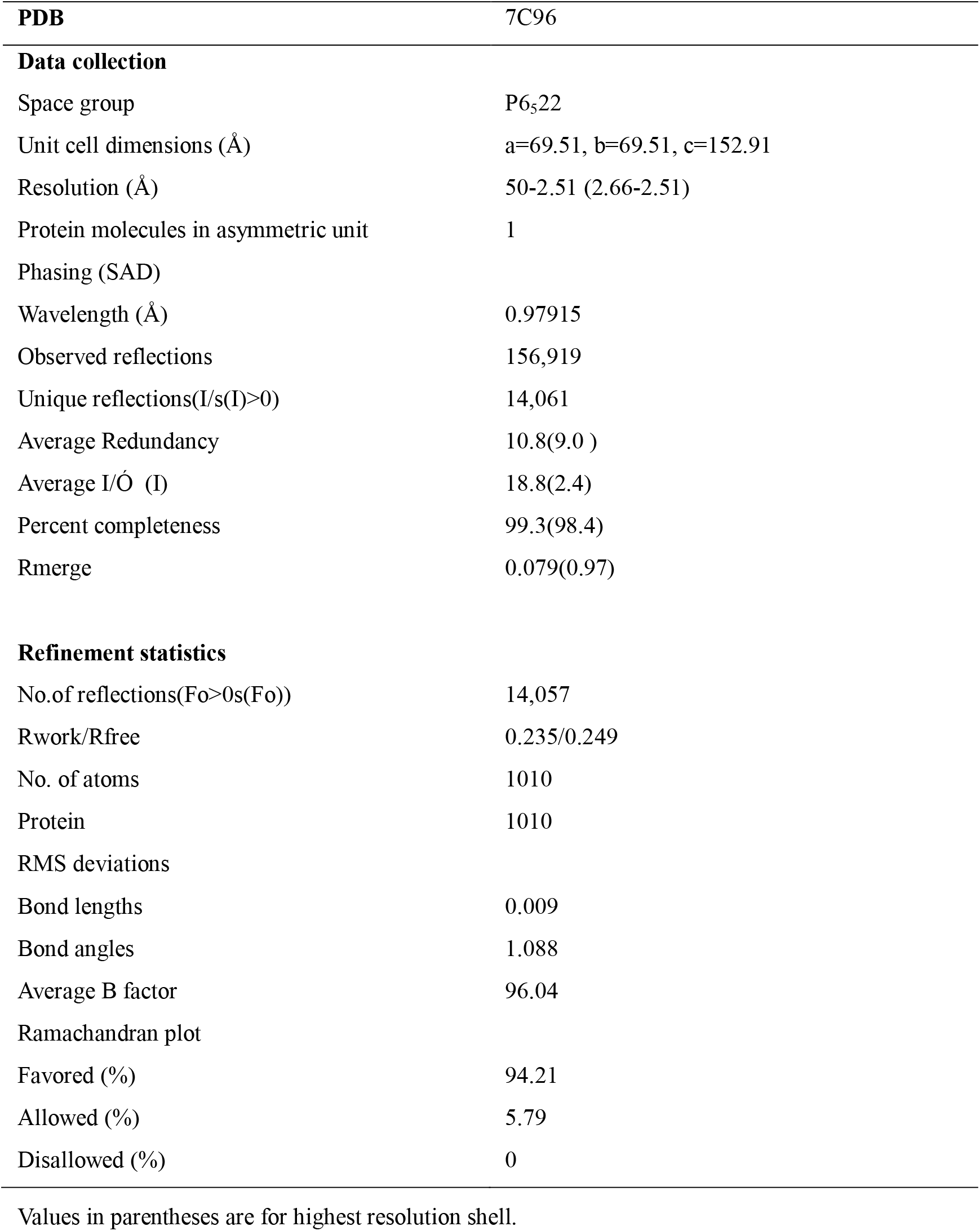
Data collection and refinement statistics.

**Table S2.**
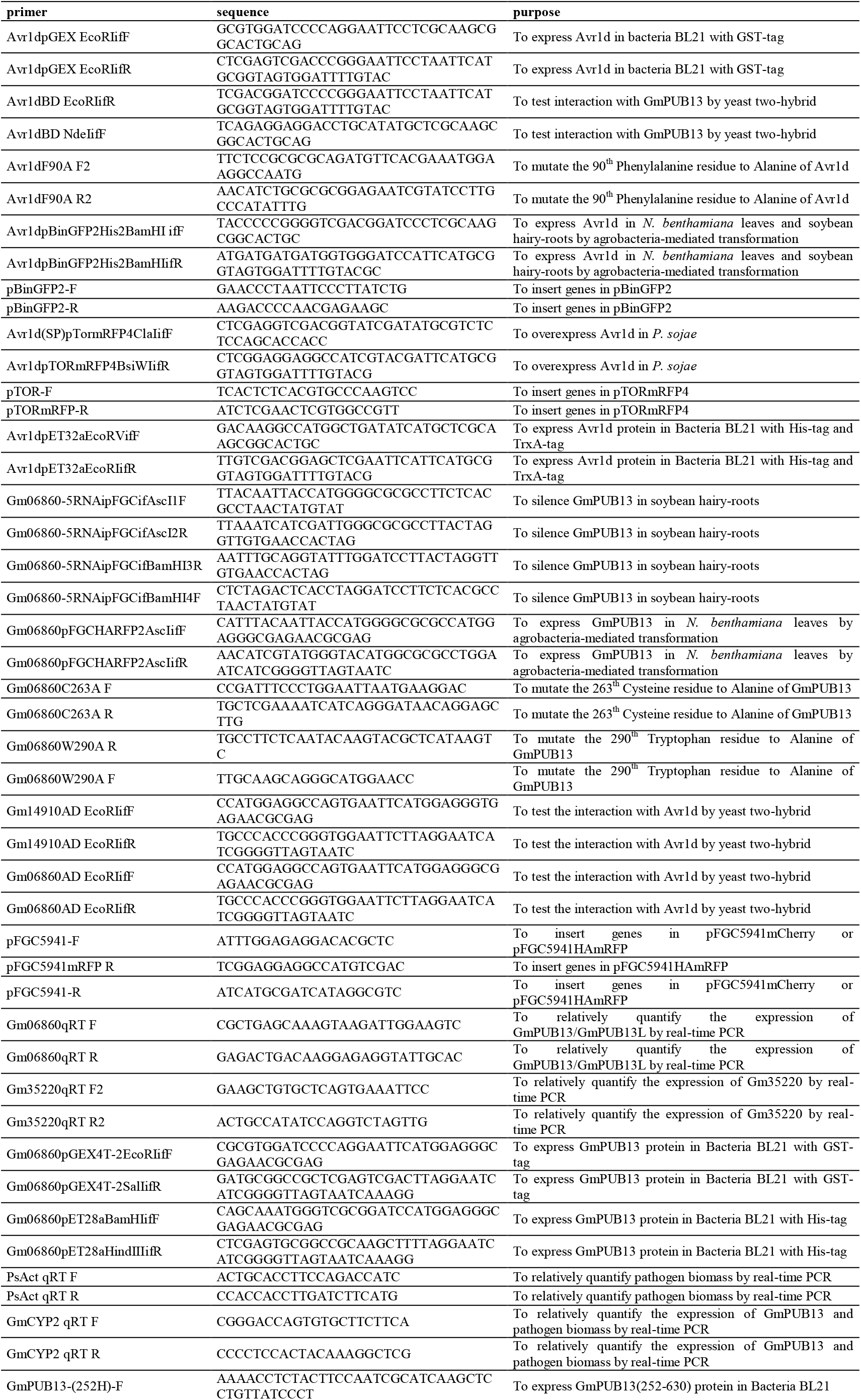
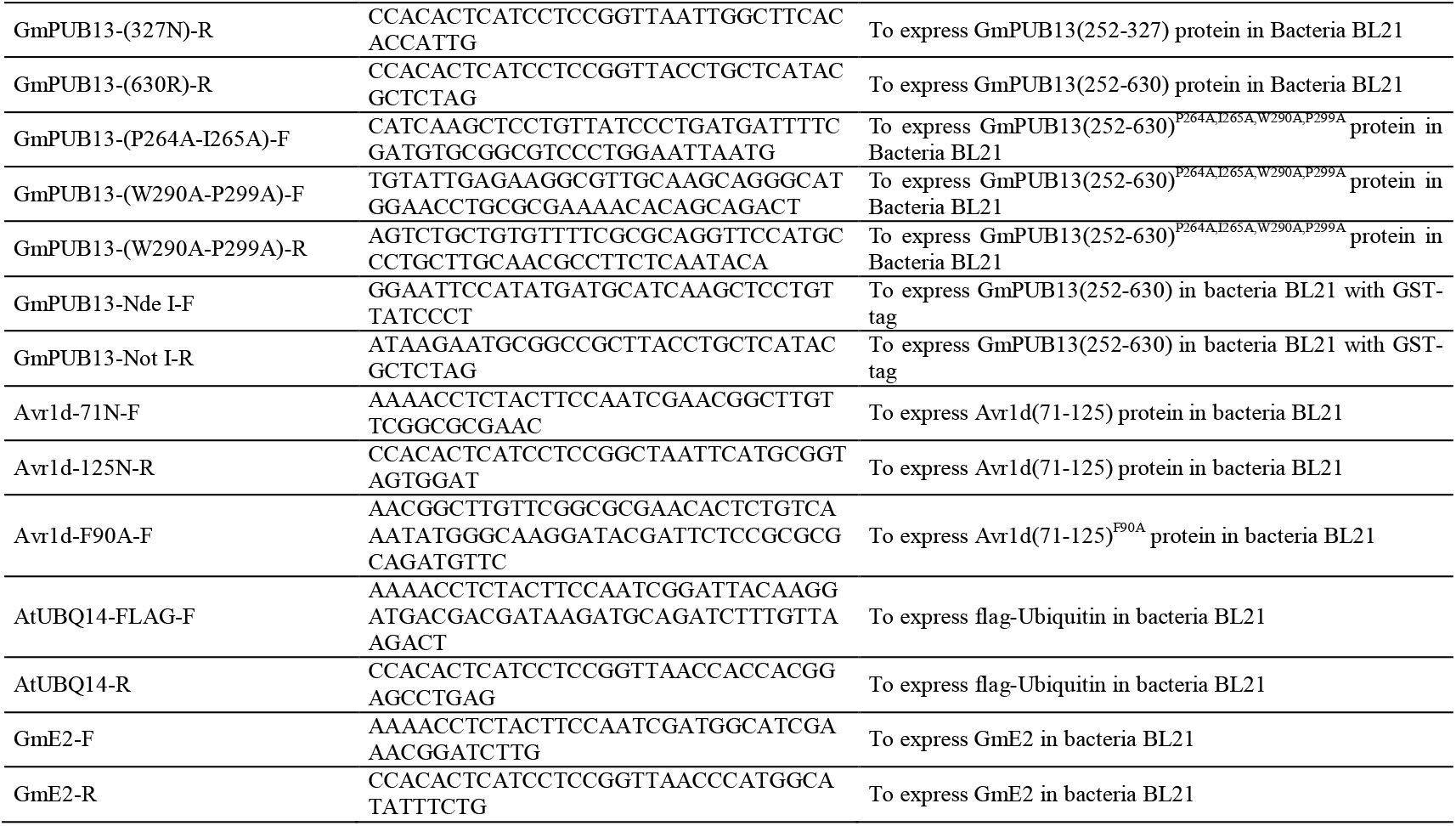
Primers used in this research.

**Table S3.**
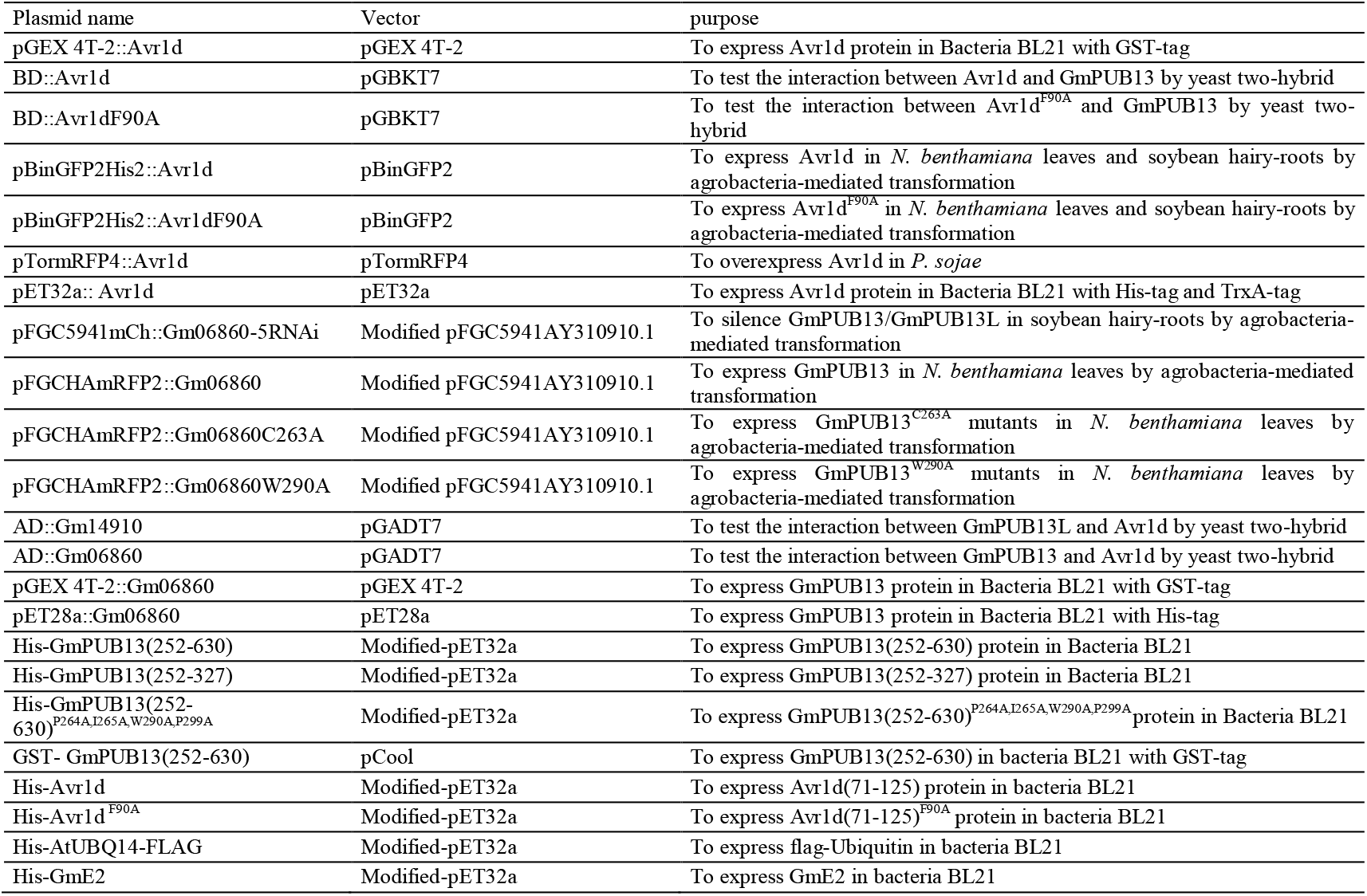
Plasmid used in this research.

